# TRIM21 licenses pancreatic acinar fate by modulating BiP phase state through VCP-dependent retrotranslocation

**DOI:** 10.64898/2026.09.29.755563

**Authors:** Yifu Zhu, Ronghua Yu, Xue Luan, Yan Huang, Quanda Li, Yuran Zhao, Huixia Zhang, Xinghua Huang, Yuxin Chen, Xiaoya Wu, Zhongjun Zhao, Jingke Ding, Jing Xue, Chong Hong, Liye Ma, Jiayu Wu, Na Wang, Jintao Xu, Wenhui Zhang, Christian Wolfrum, Tongtong Wang, Yingfang He, Jin Bai, Wenjie Ge

## Abstract

The endoplasmic reticulum (ER) chaperone BiP/GRP78 sits at the center of the proteostasis network, coordinating unfolded protein response (UPR) signaling and supporting oncogenic programs of plasticity, metastasis, immune evasion, and chemoresistance. While BiP undergoes liquid–liquid phase separation in vitro, whether mammalian cells dynamically tune this physicochemical behavior to prevent functional exhaustion or pathological condensation under sustained ER stress has remained unclear. Here, we identify the E3 ubiquitin ligase TRIM21 as a phase-state regulator that suppresses BiP’s liquid-to-solid phase separation, clearing, and preventing insoluble aggregates—thereby controlling BiP’s localization and availability to sustain the adaptive UPR under ER stress. We found that during ER stress, BiP is prone to liquid-to-solid phase separation, a transition that renders the chaperone dysfunctional. TRIM21 is recruited to BiP, where it directly polyubiquitinates BiP at specific K547/553/554/556 sites with K6-, K27-, K33-, and K63-linked chains; VCP recognizes these marks and extracts BiP from the ER lumen via ER-associated degradation (ERAD), coupling ubiquitin tagging to retrotranslocation. This preserves chaperone availability and sustains adaptive UPR—a homeostatic mechanism broadly relevant to UPR-driven diseases. In vivo, genetic ablation of *Trim21* in *Kras*^G12D/+^; *Pdx1*^Cre/+^ mice triggers aberrant BiP solidification while curtailing acinar-to-ductal transdifferentiation and tumorigenesis; notably, TRIM21 co-localizes with BiP condensates in pre-malignant human pancreatic lesions, underscoring the clinical relevance of this regulatory axis. Our findings define the TRIM21-BiP axis as a phase-control system at the interface of proteostasis and stress adaptation, offering a new framework with translational potential.

**Significance statement:** BiP phase separation is controlled by ubiquitin-dependent retrotranslocation, linking the UPR machinery to oncogenic reprogramming. We identify TRIM21 as a phase-state rheostat of the ER chaperone BiP and show that the TRIM21–BiP axis defines an unexpected interface between proteostasis and stress adaptation, revealing BiP phase behavior as a potential therapeutic target in early-stage pancreatic cancer.

## INTRODUCTION

Solid tumors are continuously subjected to intrinsic and microenvironmental insults that impose enduring stress and elicit adaptive programs essential for cancer cell survival and malignant progression (1–3). As a central hub for protein biosynthesis, modification, and trafficking, the endoplasmic reticulum (ER) maintains a proteostatic balance that is exquisitely vulnerable to oncogenic signaling, hypoxia, and nutrient deprivation. These perturbations disrupt ER function, promote the accumulation of misfolded proteins, and ultimately engage ER stress. To counteract proteotoxic damage, tumor cells have evolved sophisticated quality-control networks that restore ER homeostasis by enhancing protein folding and clearing aberrant polypeptides. Among these, ER-associated degradation (ERAD) and the unfolded protein response (UPR) constitute two core regulatory arms (4). ERAD eliminates misfolded ER substrates through the coordinated action of E3 ubiquitin ligases, retrotranslocation channels, and the AAA+ ATPase VCP/p97 (5). Concurrently, the UPR integrates stress signaling through three canonical branches—protein kinase RNA-like ER kinase (PERK), activating transcription factor 6α (ATF6α), and inositol-requiring enzyme 1 (IRE1)—which transcriptionally reinforce ERAD-associated gene expression (6).

The ER-resident Hsp70 chaperone BiP (GRP78/HSPA5) serves as the central effector of this adaptive network, governing client-protein folding and UPR homeostasis by sequestering the three UPR sensors in a latent state. To date, most studies have centered on how BiP abundance shapes tumorigenesis. Reduced BiP expression dampens PI3K/AKT and EGFR signaling, restricts pancreatic acinar-to-ductal metaplasia (ADM) (7), and suppresses *Kras*-driven tumorigenesis across multiple cancer types, largely through excessive UPR-mediated cell death (8–12). Pancreatic acinar cells function as massive protein-secretion factories, rendering their ER proteostasis network arguably the most highly burdened in the mammalian body (13). During oncogenic KRAS-driven ADM, the surging demand for protein folding imposes extreme ER stress. Classic genetic models have demonstrated this profound dependence: BiP haploinsufficiency catastrophically suppresses PI3K/AKT signaling, aborts ADM initiation, and ultimately halts KRAS-driven pancreatic tumorigenesis (7). Yet chaperone function depends not only on protein quantity but also on its intrinsic biophysical properties. Emerging evidence indicates that ER protein phase separation critically controls organelle remodeling and tumor progression (14). It has been observed that BiP inherently undergoes reversible oligomerization and spontaneous clustering within the ER (15). Indeed, recent biophysical and biochemical evidence has formalized this phenomenon, demonstrating that BiP undergoes liquid–liquid phase separation (LLPS) in vitro (16) and can be solidified by Aβ40 (17). How mammalian cells dynamically tune BiP physicochemical states to avert functional exhaustion and pathological condensation under persistent ER stress remains largely unexplored.

Here, we uncover a previously unappreciated biophysical regulatory axis, TRIM21–BiP, that couples phase-state control to ER stress adaptation and pancreatic acinar fate. We establish that the E3 ubiquitin ligase TRIM21 acts as a pivotal determinant of BiP phase behavior. Under sustained ER stress, BiP undergoes aberrant liquid-to-solid phase transition, sequestering functional BiP and thereby compromising adaptive UPR signaling.

Mechanistically, TRIM21 is recruited to BiP and catalyzes its polyubiquitination at K547/553/554/556 sites with K6-, K27-, K33-, and K63-linked chains. These modified chains are subsequently recognized by VCP to extract BiP and preserve a soluble, functionally competent chaperone pool. In *Kras*^G12D/+^; *Pdx1*^Cre/+^ mice, *Trim21* ablation triggers excessive BiP solidification, disrupts adaptive UPR homeostasis, and markedly suppresses ADM formation and pancreatic tumorigenesis. Collectively, our findings define the TRIM21–BiP axis as a critical phase-control module that bridges proteostasis and stress adaptation, offering a new framework with translational potential.

## RESULTS

### 1. Loss of TRIM21 converts physiological BiP liquid–liquid phase separation into solid-like aggregates

BiP undergoes LLPS in vitro (16), yet how this behavior is regulated in cells—and what preserves a dynamic, functional BiP pool—remains unresolved. Given that ER proteostasis is perturbed across diverse pathological states—including neurodegeneration, metabolic disease, inflammation, and cancer—identifying the molecular mechanisms that govern BiP homeostasis is of broad physiological relevance. To address this, we employed a layered, hypothesis-driven screen. Because inflammatory and chemically-induced ER stress represents a shared trigger of BiP dysfunction across tissues (18), we reasoned that stress-responsive E3 ubiquitin ligases would constitute the most physiologically relevant regulators of the BiP network. We therefore first interrogated our in-house RNA-seq dataset of caerulein-induced pancreatitis—a well-characterized, tissue-level model of acute inflammatory ER stress—to capture stress-induced E3 ubiquitin ligases. To prioritize candidates most likely to act on BiP in a clinically meaningful context, we intersected this stress-responsive genes with three additional criteria: (i) ER stress–associated genes; (ii) E3 ubiquitin ligases ome (19); and (iii) predicted BiP interactors (20). This intersection uniquely nominated TRIM21 (Figure 1A). A second convergence step—merging predicted TRIM21 interactors, previously reported (21) and our own TRIM21 Co-IP mass spectrometry datasets, and the UPR pathway—further positioned TRIM21 as a BiP-associated E3 ligase at the nexus of ER proteostasis and stress adaptation (Figure 1B). Consistent with a physical interaction, endogenous BiP and TRIM21 colocalized in perinuclear puncta under basal conditions, and this colocalization was markedly enhanced by two mechanistically distinct ER-stress inducers—thapsigargin, which depletes ER calcium stores (22), and tunicamycin, which inhibits N-linked glycosylation (23)—thereby corroborating the physiological relevance of their association (Figure 1C and S1A). Notably, TRIM21 deletion in PANC-1 and MEF cells produced significantly enlarged condensates (Figure 1D, 1E, S1B and S1C). Because TRIM21 reportedly promotes the ubiquitin-mediated degradation of G3BP1 (24), a core stress-granule component, we asked whether BiP condensation is linked to stress granules. Using MEFs stably co-expressing Cherry-BiP and GFP-G3BP1, we found that BiP condensates do not colocalize with G3BP1 under either inducer, indicating that TRIM21-driven BiP condensation occurs independently of the canonical stress-granule pathway (Figure S1D).

**Figure 1.**
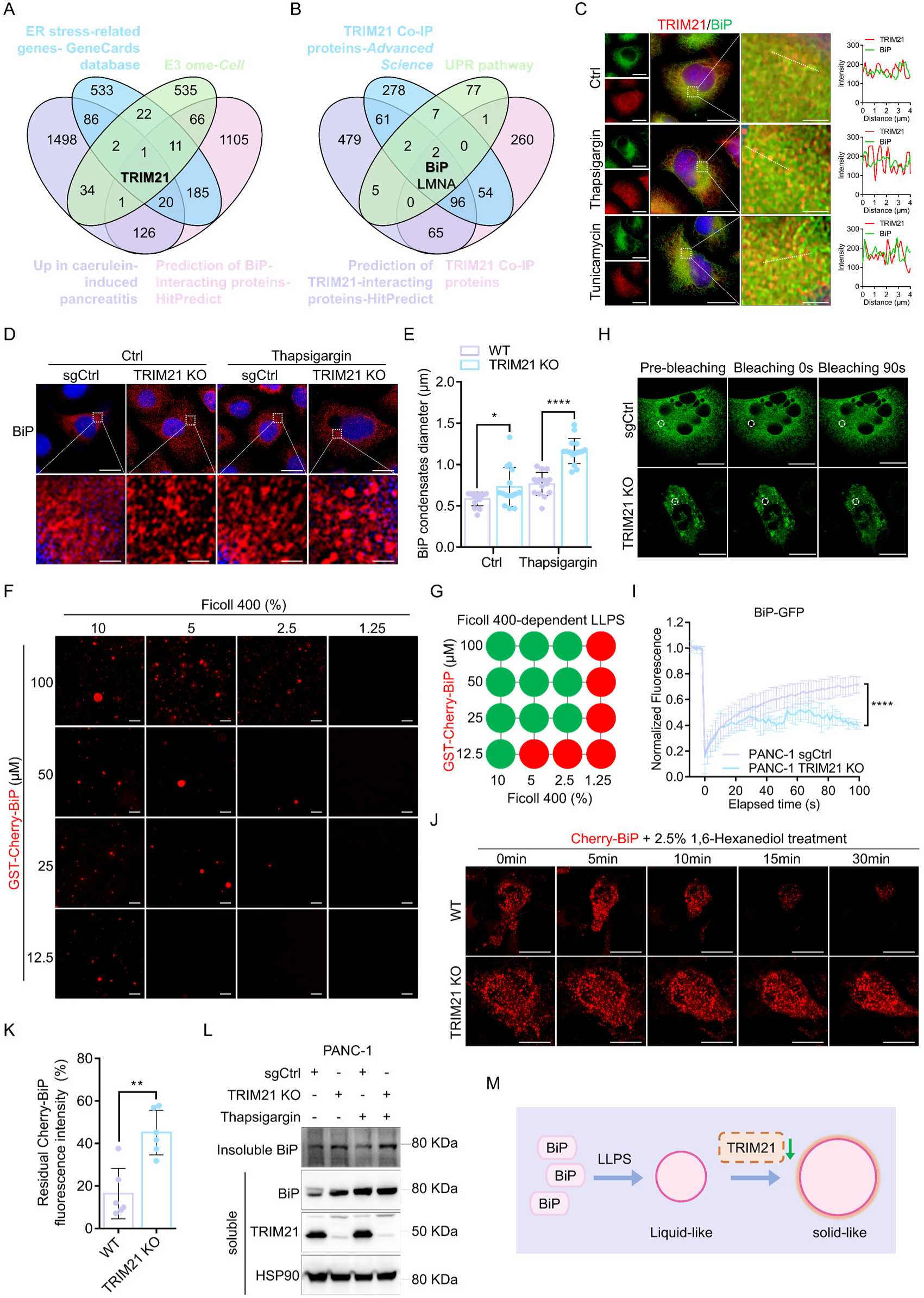
Loss of TRIM21 converts physiological BiP liquid-liquid phase separation into solid-like aggregates. (A) Venn diagram showing the overlap of genes among four independent datasets: genes upregulated in caerulein-induced pancreatitis, ER stress-related genes from the GeneCards database, E3 ubiquitin ligase ome, and HitPredict-predicted BiP-interacting proteins. (B) Venn diagram showing the overlap among HitPredict-predicted TRIM21-interacting proteins, proteins identified in two independent TRIM21 Co-IP experiments, and the UPR pathway. (C) Immunofluorescence analysis of BiP and TRIM21 in PANC-1 cells treated with DMSO (Ctrl), 1 μM Thapsigargin, or 5 μg/mL Tunicamycin for 4 hours. Scale bar, 20 μm and 2 μm. (D and E) Immunofluorescence analysis of BiP in PANC-1 sgCtrl and TRIM21 KO cells treated with DMSO (Ctrl) or 1 μM Thapsigargin for 4 hours. Scale bar, 20 μm and 2 μm. Diameter of BiP condensates in (D), presented as mean ± SD for 15 cells of 3 independent experiments (E). *: *P*<0.05, ****: *P*<0.0001 (unpaired Student’s two-tailed t-test). (F and G) Phase-separated droplets of GST-Cherry-BiP imaged in a Ficoll-400-based crowding environment, with concentration-dependent effects documented through representative microscopy (F) and summary of phase separation behaviors of BiP at the indicated concentrations (G). Scale bar, 20 µm. (H) PANC-1 sgCtrl and TRIM21 KO cells transduced with lenti-EFs-BiP-GFP plasmid were subjected to fluorescence recovery after photo-bleaching (FRAP) using a Leica STELLARIS 5 confocal microscope (488 nm channel). Time-lapse imaging captured BiP-GFP condensates recovery. Scale bar, 20 µm. (I) Quantification of BiP-GFP condensates FRAP in PANC-1 sgCtrl and TRIM21 KO cells. Data were presented as mean ± SD of 4 bleached areas. ****: *P*<0.0001 (two-way ANOVA with Tukey’s multiple-comparisons test). (J) Confocal time-lapse imaging revealed that 1,6-hexanediol (2.5%) promotes the dissociation of Cherry-BiP condensates in wild-type (WT) and TRIM21 KO MEF cells. Scale bar, 20 µm. (K) Quantification of residual Cherry-BiP fluorescence intensity in (J). Data were mean ± SD of 6 independent experiments. **: *P*<0.01 (unpaired Student’s two-tailed t-test). (L) Relative protein levels of BiP in the soluble and insoluble fractions of PANC-1 sgCtrl and TRIM21 KO cells treated with DMSO (Ctrl) or 1 μM Thapsigargin for 4 hours. HSP90 was used as a soluble fraction loading control. (M) Schematic model depicting that BiP can undergo LLPS; in the absence of TRIM21, BiP instead undergoes a liquid-to-solid phase transition.

We next asked how these condensates are regulated, beginning with the intrinsic phase-separation capacity of BiP. Prokaryotically expressed GST-Cherry-BiP robustly formed droplets at low concentrations of the crowding agent Ficoll 400 in vitro (Figure 1F and 1G), whereas GST-Cherry alone required substantially higher Ficoll concentrations (Figure S1E and S1F). In cells, fluorescence recovery after photobleaching (FRAP) showed that BiP-GFP condensates recovered rapidly in controls but displayed markedly impaired recovery in TRIM21-KO cells, indicating a loss of liquid-like dynamics (Figure 1H and 1I). Accordingly, the droplet-disrupting agent 1,6-hexanediol dissolved BiP condensates efficiently in wild-type (WT) but only partially in TRIM21 KO cells (Figure 1J and 1K), and biochemical fractionation revealed that ER stress drove BiP into an insoluble, solid-like fraction preferentially in TRIM21 KO cells (Figure 1L). Collectively, these data establish that BiP forms physiological liquid condensates during ER stress and that TRIM21 maintains this reversible liquid state; in its absence, BiP undergoes an aberrant liquid-to-solid transition that sequesters the chaperone (Figure 1M).

### 2. TRIM21 ubiquitinates BiP at SBDα/lid lysines, licensing its proteasomal turnover and restoring liquid-phase behavior

After establishing that TRIM21 governs the phase behavior of the ER chaperone BiP, we set out to define the molecular basis underlying this regulation. Co-expressing Flag-TRIM21 with HA-tagged BiP and the UPR sensors PERK, IRE1α, ATF6, ATF4 in cells, we found that TRIM21 associated selectively with BiP (Figure S2A) but with none of the UPR sensors under identical conditions (Figure S2B–S2E). This selectivity was recapitulated in vitro, as recombinant TRIM21 pulled down BiP in GST pulldown assays (Figure 2A). Truncation analysis of HA-tagged BiP deletion mutants narrowed the TRIM21-binding interface to the substrate-binding domain (SBD; residues 420–552): all SBD-containing variants co-precipitated with TRIM21, whereas SBD-lacking constructs failed to do so (Figures 2B and 2C). Reciprocally, domain dissection of TRIM21 mapped its BiP-binding activity to the C-terminal PRY-SPRY domain (Figures S2F and S2G), a module canonically dedicated to substrate recognition. In consistent silico docking of the TRIM21 dimer (PDB: 2IWG) onto BiP (PDB: 6ASY) predicted an interface involving TRIM21 residue Y296 and BiP residues P438/T445/K446 (Figure S2H). Thus, TRIM21 engages BiP through a site-specific PRY-SPRY–SBD interaction.

**Figure 2.**
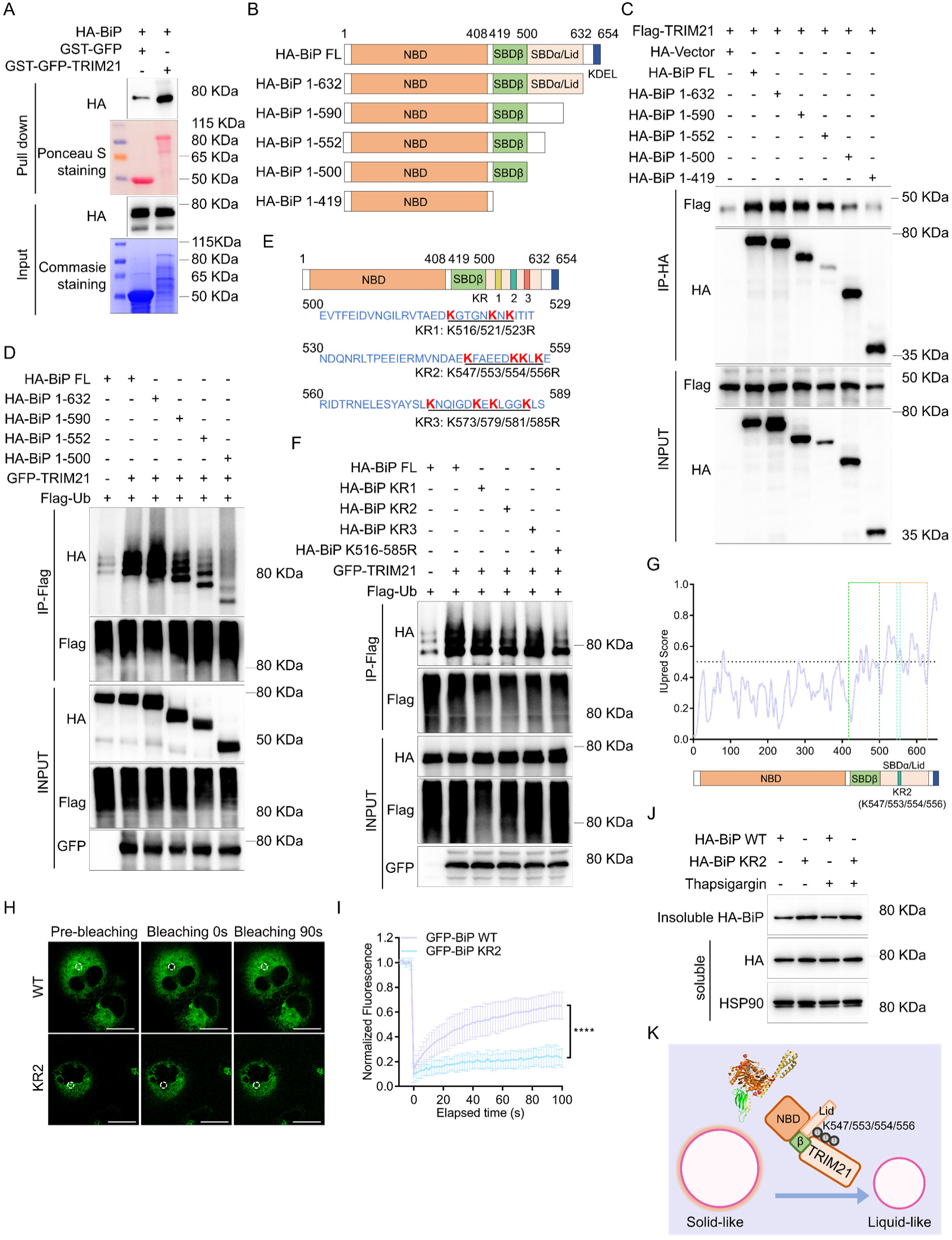
TRIM21 binds the BiP substrate-binding domain (SBD, residues 420-552) and ubiquitinates SBDα/lid residues K547, K553, K554, and K556. (A) Purified recombinant GST-GFP or GST-GFP-TRIM21 was incubated with recombinant HA-BiP; bound BiP was analyzed by Western blotting. (B) Schematic of the cloning strategy used to generate HA-BiP truncation mutants. (C) Co-immunoprecipitation (Co-IP) assays in HEK-293T cells co-transfected with Flag-TRIM21 and either full-length BiP or the indicated BiP truncation mutants. Immunoprecipitates (top) and inputs (bottom) were analyzed by Western blotting against anti-HA or anti-Flag. (D) Ubiquitination assays in HEK-293T cells co-transfected with full-length HA-BiP or the indicated truncation mutants, along with Flag-Ub and GFP-TRIM21. Samples recovered on Flag-beads (top) and inputs (bottom) were analyzed by Western blotting against anti-HA, anti-Flag, and anti-GFP as indicated. (E) Schematic of HA-BiP lysine-substitution mutants: KR1 (K516/K521/K523R), KR2 (K547/K553/K554/K556R), KR3 (K573/K579/K581/K585R), and K516-585R. (F) Ubiquitination assays were performed in HEK-293T cells as in (D), after co-transfection with full-length HA-BiP or the indicated KR mutants, in the presence or absence of GFP-TRIM21. (G) Intrinsic disorder regions of BiP predicted by IUPred3 (scores >0.5 indicate disorder). The ubiquitinated residues (KR2: K547/K553/K554/K556) fall within these disordered regions. (H) PANC-1 cells transduced with lenti-EFs-BiP-WT-GFP or lenti-EFs-BiP-KR2-GFP were subjected to FRAP analysis on a Leica STELLARIS 5 confocal microscope (488 nm channel). Time-lapse imaging captured BiP-GFP condensate recovery. Scale bar, 20 μm. (I) Quantification of FRAP recovery of BiP WT-GFP and KR2-GFP condensates in PANC-1 cells, presented as mean ± SD of 4 bleached areas. *: *P*<0.05 (two-way ANOVA with Tukey’s multiple-comparisons test). (J) Relative protein levels of HA-BiP WT and HA-BiP KR2 in the soluble and insoluble fractions of HEK-293T cells treated with DMSO (Ctrl) or 1 μM Thapsigargin for 4 hours. HSP90 was used as a soluble fraction loading control. (K) Schematic model showing that TRIM21 binds to the BiP SBD and ubiquitinates residues K547, K553, K554, and K556 within the SBDα/lid subdomain.

We next asked whether this recognition licenses BiP ubiquitination. Flag-Ub-based assays in HEK-293T cells showed that TRIM21 co-expression selectively promoted BiP ubiquitination (Figure S3A), whereas knockdown of endogenous TRIM21 reduced it (Figure S3B). Functionally, TRIM21 knockdown elevated BiP protein levels (Figure S3C), while TRIM21 overexpression reduced them—an effect reversed by the proteasome inhibitor MG132 (Figure S3D), implicating proteasomal degradation. Consistently, cycloheximide (CHX) chase experiments showed that TRIM21 knockdown stabilized endogenous BiP (shCtrl: 6 hours vs shTRIM21: >6 hours; Figures S3E and S3F), whereas ectopic TRIM21 destabilized exogenous HA-BiP (Flag-Vector: >8 hours vs Flag-TRIM21: 6 hours; Figures S3G and S3H). We conclude that TRIM21-mediated ubiquitination routes BiP to the proteasome.

To identify the modified lysines, we profiled BiP C-terminal truncation mutants and localized the principal ubiquitination sites to residues 500–590aa (Figure 2D), a stretch bearing eleven lysines. Mutational analysis revealed that, uniquely among the mutants tested, KR2 (K547/553/554/556R) reduced BiP ubiquitination comparably to the K547-585R mutant, identifying K547/K553/K554/K556 as the primary TRIM21 substrates (Figure 2E and 2F). TRIM21 overexpression did not alter HA-BiP-KR2 levels (Figure S3I), indicating that proteasomal degradation was blocked; accordingly, KR2 accumulated in CHX-treated cells (HA-BiP WT: 8 hours vs HA-BiP KR2: >8 hours; Figures S3J and S3K). Notably, KR2 did not affect BiP–TRIM21 interaction but slightly diminished BiP binding to ATF6 (Figures S3L and S3M). Using linkage-specific Flag-Ub mutants, we found that TRIM21 promotes K6-, K27-, K33-, and K63-linked polyubiquitination of BiP (Figure S3N). In summary, TRIM21 acts as a negative regulator of BiP by promoting its K6-, K27-, K33-, and K63-polyubiquitination and proteasomal degradation.

These ubiquitination sites (K547/553/554/556) map to an intrinsically disordered region (IDR) of BiP (Figure 2G), where polyubiquitin chains can remodel both steric and electrostatic surface properties (25). Because polyubiquitin serves as a multivalent platform for ubiquitin-binding/associated domains (UBDs/UBAs) within condensates and thereby dynamically regulates LLPS (26), we asked whether BiP ubiquitination governs its phase behavior. In PANC-1 cells stably expressing BiP-WT-GFP or BiP-KR2-GFP, KR2 foci displayed a markedly attenuated FRAP profile, indicating that ubiquitin modification regulates BiP’s liquid-to-solid transition (Figures 2H and 2I). Consistently, wild-type BiP remained predominantly in the soluble fraction whereas KR2 mutant partitioned toward the insoluble pool (Figure 2J). Given that UBQLN2 couples ubiquitinated clients to the proteasome (27), we examined whether proteasomes engage BiP condensates. PSMB5 did not fully co-localize with BiP condensates, yet their proximity was significantly reduced in TRIM21 KO cells (Figure S3O). Collectively, these data demonstrate that TRIM21-mediated K6-, K27-, K33-, and K63-polyubiquitination of the SBDα/lid residues K547/553/554/556 within the phase-regulatory IDR functions as a biophysical shield, which counteracts aberrant liquid-to-solid transition of BiP (Figure 2K).

### 3. Ubiquitinated BiP is retrotranslocated to the cytosol and degraded via the VCP-dependent ERAD system

Having established that TRIM21 ubiquitinates BiP at its SBDα/lid region, we next asked how this ER-luminal chaperone could serve as a substrate for the cytosolic E3 ligase TRIM21. Because BiP relocalizes from the ER lumen to the nucleus (28, 29) and plasma membrane (30, 31) under stress, we reasoned that stress-induced BiP may transiently become accessible to cytosolic TRIM21 and sought to define the site and mechanism of this engagement. To determine whether cytosolic TRIM21 encounters BiP at the ER surface rather than after cytoplasmic leakage, we first examined the subcellular topology of BiP condensates under distinct stress regimes. In TRIM21 KO MEFs, thapsigargin-induced BiP condensates were enveloped by SEC61β, an ER membrane translocon component (Figure S4A). Likewise, MG132-induced BiP condensates remained confined to the ER and were co-enveloped by SEC61β (Figure S4A), indicating that the observed association reflects ER-resident localization rather than release of BiP into the cytosol. To directly visualize BiP-TRIM21 proximity, we performed bimolecular fluorescence complementation (BiFC) using split GFP. Luminal Cherry-BiP-GFP1-10 and cytosolic GFP11-TRIM21 reconstituted a functional fluorophore, producing a robust near ER-localized green signal (Figure S4C and S4D). This in situ reconstitution demonstrates that cytosolic TRIM21 engages luminal BiP across the ER membrane, revealing an ER-surface-based ubiquitination platform that expands the conventional view of TRIM21 as a purely cytosolic E3 ligase.

Given that ER-luminal proteins are cleared primarily through ERAD, where they are ubiquitinated and degraded by the proteasome. Upon comparing the proteins co-immunoprecipitated with TRIM21 in PANC-1 cells to known ERAD components, we found that only BiP and the retrotranslocation ATPase VCP/p97 were present (Figure 3A). In thapsigargin-treated MEFs, BiP, TRIM21, VCP, and SEC61β colocalized (Figure 3B).

**Figure 3.**
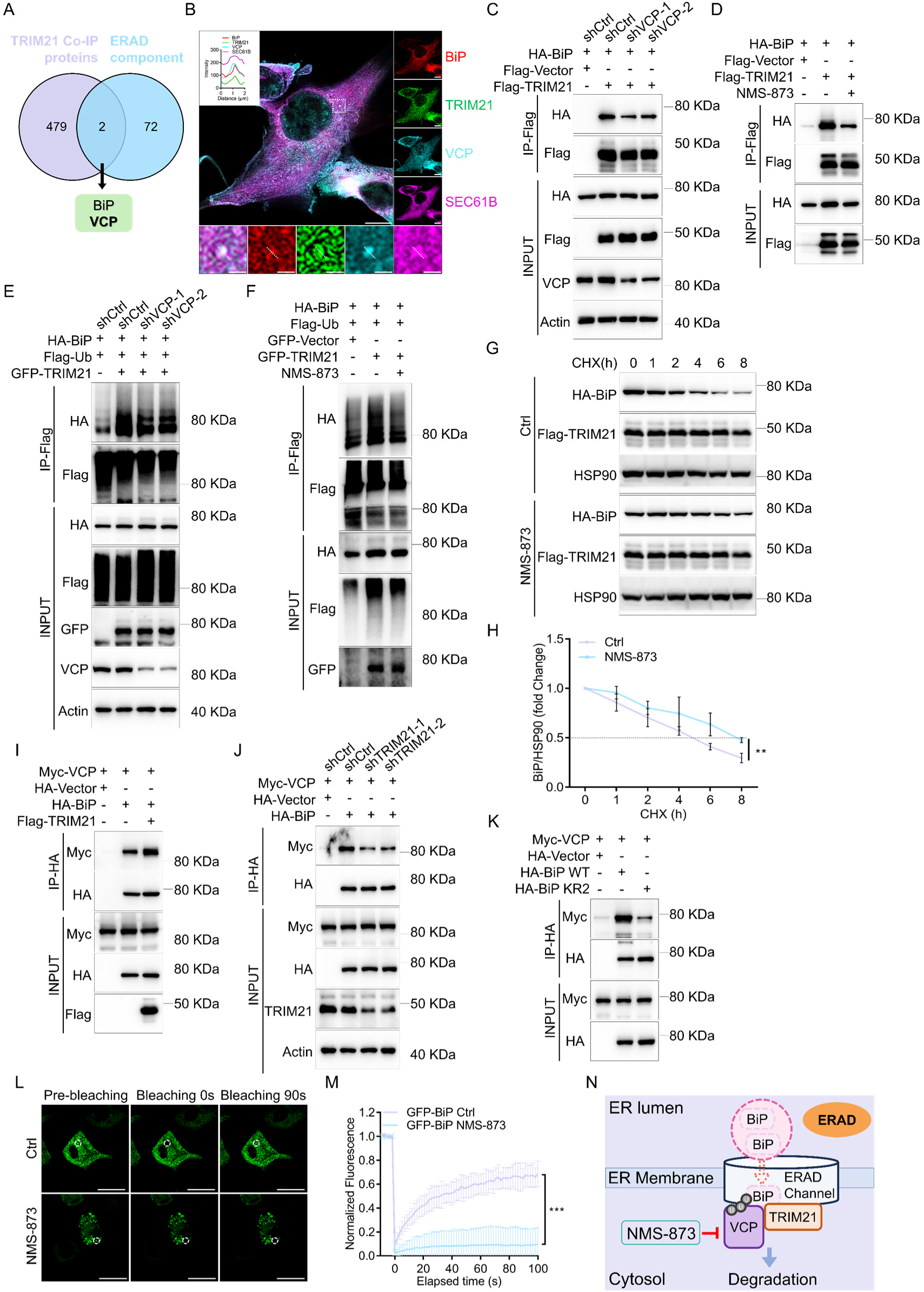
Ubiquitinated BiP is retrotranslocated and degraded via the VCP-dependent ERAD system. (A) Venn diagram of TRIM21-binding proteins identified by TRIM21 co-IP in PANC-1 cells, compared with ERAD component genes. (B) Immunofluorescence analysis of BiP, TRIM21, VCP and SEC61β in WT MEF cells treated with 1 μM Thapsigargin for 4 hours. Scale bar, 20 μm and 2 μm. (C) Co-IP assays in HEK-293T shCtrl, shVCP-1, and shVCP-2 cells co-transfected with HA-BiP and Flag-TRIM21. Immunoprecipitates (top) and inputs (bottom) were analyzed by Western blotting against anti-HA, anti-Flag, anti-VCP, and anti-Actin. (D) Co-IP assays in HEK-293T cells co-transfected with HA-BiP and Flag-TRIM21, treated with DMSO (Ctrl) or 2 μM NMS-873 for 4 hours. Immunoprecipitates (top) and inputs (bottom) were analyzed by Western blotting against anti-HA and anti-Flag. (E) Ubiquitination assays in HEK-293T shCtrl, shVCP-1, and shVCP-2 cells. Samples recovered on Flag-beads (top) and inputs (bottom) were analyzed by blotting against anti-HA, anti-Flag, anti-GFP, anti-VCP, and anti-Actin as indicated. (F) Ubiquitination assays in HEK-293T cells treated with DMSO (Ctrl) or 2 μM NMS-873 for 4 hours. Samples recovered on Flag-beads (top) and inputs (bottom) were analyzed by blotting against anti-HA, anti-Flag, and anti-GFP. (G and H) Western blot analysis of HA-BiP levels in HEK-293T cells treated with cycloheximide (CHX; 100 μg/mL) with DMSO (Ctrl) or 2 μM NMS-873 for 4 hours; blots were probed against anti-HA, anti-TRIM21, and anti-HSP90 (loading control). Western blot analysis of HA-BiP levels in 100 μg/mL CHX-treated HEK-293T cells with or without 2 μM NMS-873 for 4 hours (G). Densitometric quantification of HA-BiP levels from (G), presented as mean ± SD of 3 independent experiments (H). **: *P*<0.01 (two-way ANOVA with Tukey’s multiple-comparisons test). (I) Co-IP assays in HEK-293T cells co-transfected with Myc-VCP and HA-BiP, with or without Flag-TRIM21. Immunoprecipitates (top) and inputs (bottom) were analyzed by Western blotting against anti-Myc, anti-HA, and anti-Flag. (J) Co-IP assays in HEK-293T shCtrl, shTRIM21-1, and shTRIM21-2 cells co-transfected with Myc-VCP and HA-BiP. Immunoprecipitates (top) and inputs (bottom) were analyzed by Western blotting against anti-Myc, anti-HA, anti-Flag, anti-TRIM21, and anti-Actin. (K) Co-IP assays in HEK-293T cells co-transfected with Myc-VCP and HA-BiP WT or HA-BiP KR2 mutant. Immunoprecipitates (top) and inputs (bottom) were analyzed by Western blotting against anti-Myc or anti-HA. (L) PANC-1 cells transduced with lenti-EFs-BiP-WT-GFP treated with DMSO (Ctrl) or 2 μM NMS-873 for 4 hours, were subjected to FRAP analysis on a Leica STELLARIS 5 confocal microscope (488 nm channel). Time-lapse imaging captured BiP-GFP condensate recovery. Scale bar, 20 μm. (M) Quantification of FRAP recovery of BiP WT-GFP condensates in PANC-1 cells treated with DMSO (Ctrl) or 2 μM NMS-873 for 4 hours, presented as mean ± SD of 4-5 bleached areas. ***: *P*<0.001 (two-way ANOVA with Tukey’s multiple-comparisons test). (N) Schematic model showing that VCP, via ERAD, facilitates TRIM21-dependent ubiquitination of ER-luminal BiP, extracting BiP from the ER lumen into the cytosol, where TRIM21 acts as the E3 ligase.

Notably, VCP depletion or pharmacological inhibition with NMS-873 (32) reduced TRIM21–BiP interaction, suppressed BiP ubiquitination, and stabilized BiP in CHX-chase experiments (Figure 3C–3H). Likewise, loss of ERAD channels SEC61β, HRD1 (SYVN1), or DERL1 impaired both TRIM21– BiP interaction and TRIM21-mediated BiP ubiquitination (Figure S5A–S5I), suggesting that BiP must traverse an ERAD transporter to become a TRIM21 substrate.

BiP truncation analyses showed that VCP and SEC61β both bind SBD-containing BiP forms, mapping to residues 500–552 and 420–500, respectively (Figure S6A). TRIM21 promoted VCP recruitment to BiP, as overexpression enhanced and depletion weakened the VCP–BiP interaction (Figure 3I and 3J). The BiP KR2 mutant retained SEC61β binding yet exhibited reduced VCP association, indicating that ubiquitination at K547/K553/K554/K556 is required for VCP recognition (Figure 3K and S6B). Notably, NMS-873 weakened SEC61β–BiP interaction (Figure S6C), and SEC61β overexpression diminished both TRIM21–BiP and VCP–BiP interactions without affecting VCP–TRIM21 binding (Figure S6D–S6F). VCP overexpression did not alter BiP–SEC61β interaction (Figure S6G), whereas BiP overexpression specifically enhanced VCP–SEC61β binding (Figure S6H). Importantly, TRIM21 did not bind to SEC61β (Figure S6I), and TRIM21 overexpression impaired BiP–SEC61β binding (Figure S6J). SEC61β overexpression also reduced BiP ubiquitination (Figure S6K). Collectively, these data place TRIM21, VCP, and SEC61β in a cooperative yet competitive network at the SBD docking site.

VCP/p97 is an AAA+ ATPase that acts as the principal mechanical driver of ERAD, extracting polyubiquitinated clients from the ER (33). Beyond this canonical retrotranslocation role, VCP also mediates the disassembly of cytoplasmic condensates: upon heat shock, ubiquitination of the core stress-granule protein G3BP1 recruits the ER-associated adaptor FAF2, which delivers G3BP1 to VCP for extraction, driving the granule’s interaction network below the percolation threshold for phase separation (34). Whether VCP exerts a similarly remote influence on the phase behavior of ER-luminal chaperones remains unexplored. Given that TRIM21 both appends polyubiquitin chains to clients and recruits the VCP motor, and that the abundant luminal BiP forms dynamic condensates in the ER under stress, we hypothesized that TRIM21-mediated polyubiquitination might couple the VCP retrotranslocation motor to the physical dissolution of BiP condensates.

Accordingly, FRAP analyses revealed that VCP depletion or NMS-873 treatment markedly reduced recovery of BiP-GFP condensates in PANC-1 cells (Figure 3L, 3M, S6L and S6M), indicating that VCP-dependent extraction of ubiquitinated BiP prevents solid-like accumulation and preserves its liquid-like behavior. Together, these findings define a retrotranslocation-coupled degradation pathway in which the cytosolic E3 ligase TRIM21 recognizes ubiquitinated BiP emerging from the ERAD channel and cooperates with VCP to extract it for proteasomal clearance (Figure 3N).

### 4. TRIM21 deficiency promotes BiP condensate formation and ATF6 nuclear import

Having established that TRIM21 degrades ubiquitinated BiP through the VCP-dependent ERAD pathway, we asked whether this regulation shapes the direction of the UPR. Since BiP sequestration governs activation of the three UPR arms, we reasoned that TRIM21-dependent clearance of aberrant BiP determines UPR output. Indeed, modulating TRIM21 levels produced corresponding changes in CHOP and sXBP1 mRNA/protein and eIF2α-S51 phosphorylation, the key downstream effectors of the ATF6, IRE1α, and PERK branches, respectively (35) (Figure 4A and S7A–S7C). Notably, ATF4, another PERK mediator, remained unchanged. TRIM21 deficiency modulated UPR activation in an ER stress intensity–dependent manner. After 4-hours thapsigargin treatment, TRIM21 depletion enhanced sXBP1 and CHOP activation without affecting viability (sgCtrl 91.33±5.56% vs KO-1 85.14±2.13% vs KO-2 87.59±3.16%, n.s.), consistent with an adaptive UPR. By contrast, after 24-hours, sXBP1 and CHOP were robustly activated irrespective of TRIM21 status, with reduced viability (70.95±0.81% vs 65.09±1.42% vs 66.71±0.33%, P<0.05), reflecting a terminal UPR (Figure S7D and S7E).

**Figure 4.**
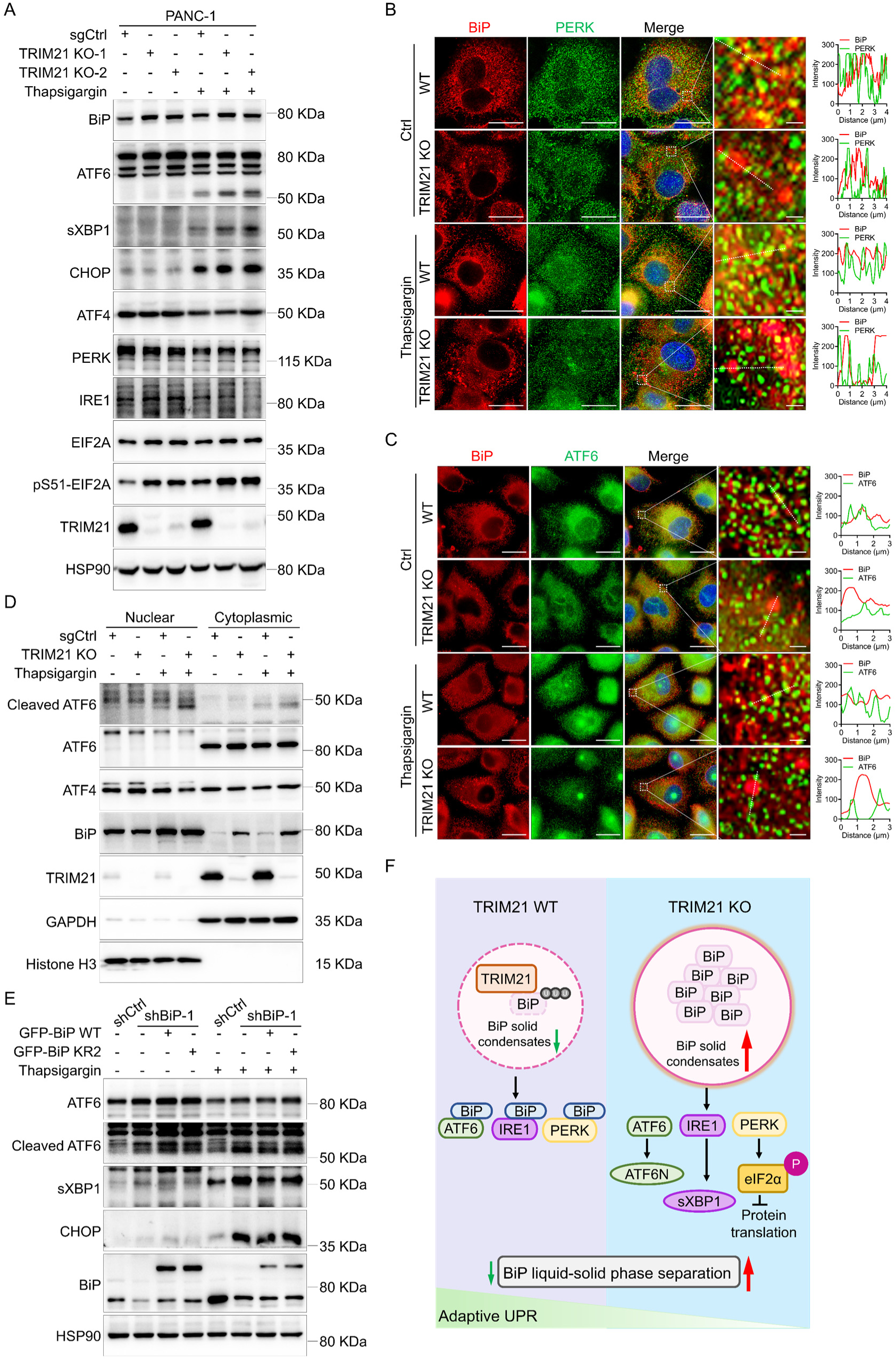
TRIM21 deficiency promotes BiP condensate formation and ATF6 nuclear import. (A) Protein levels of BiP, ATF6, sXBP1, CHOP, ATF4, PERK, IRE1, EIF2A, pS51-EIF2A, and TRIM21 in PANC-1 sgCtrl and TRIM21 KO cells treated with DMSO (Ctrl) or 1 μM Thapsigargin for 4 hours. (B and C) Immunofluorescence analysis of BiP and PERK (B) or ATF6 (C) in WT MEF and TRIM21 KO MEF cells treated with DMSO (Ctrl) or 1 μM Thapsigargin for 4 hours. Scale bar, 20 μm and 1 μm. (D) Relative protein levels of cleaved ATF6, ATF6, ATF4, BiP and TRIM21 in nuclear and cytoplasmic fractions, as determined by cell fractionation in PANC-1 sgCtrl and TRIM21 KO cells treated with DMSO (Ctrl) or 1 μM Thapsigargin for 4 hours. GAPDH and histone H3 served as loading controls for the cytoplasmic and nuclear fractions, respectively. (E) Protein levels of ATF6, cleaved ATF6, sXBP1, CHOP, BiP, and GFP-BiP in shCtrl, shBiP-1 and shBiP-1 PANC-1 cells stably complemented with BiP-WT-GFP or BiP-KR2-GFP, treated with DMSO (Ctrl) or 1 μM Thapsigargin for 4 hours. (F) Schematic model depicting that TRIM21 prevents BiP liquid-to-solid phase separation and promotes the degradation of BiP solid condensates, thereby maintaining adaptive UPR homeostasis.

Loss of TRIM21-dependent control over BiP’s physical state has functional consequences for UPR signaling. In wild-type MEFs, BiP sequesters the sensors PERK and ATF6 in an inactive state. Upon TRIM21 loss, however, aberrant BiP condensation led to sequestration failure, releasing these sensors: immunofluorescence revealed reduced colocalization of BiP condensates with both PERK and ATF6 in TRIM21 KO MEFs (Figure 4B and 4C). This dissociation unleashed downstream signaling, driving robust ATF6 nuclear translocation, as confirmed by imaging and biochemical fractionation (Figure 4C, 4D and S7F).

To establish causality, we reconstituted BiP-knockdown PANC-1 cells with BiP-WT-GFP or the KR2 ubiquitination-resistant mutant. In thapsigargin-treated cells, BiP-WT restored a balanced UPR (reduced sXBP1, CHOP, and cleaved ATF6), whereas BiP-KR2 failed to suppress these outputs and recapitulated the TRIM21 KO phenotype (Figure 4E). Likewise, VCP depletion or NMS-873 inhibition elevated insoluble BiP, cleaved ATF6, sXBP1, and CHOP (Figure S7G and S7H). Collectively, these data establish TRIM21 as a phase-state regulator that prevents dysfunctional BiP condensate formation, preserving chaperone availability to sequester ER stress sensors and safeguard adaptive UPR homeostasis (Figure 4F).

### 5. Ablation of *Trim21* in pancreatic cells inhibits PDAC tumorigenesis in KC mice

Having established that TRIM21 restrains BiP solid-phase condensation and sustains an adaptive UPR program, we asked whether this regulation impacts pancreatic tumorigenesis. Pancreatic acinar cells exhibit exceptionally high ER-dependent secretory activity, imposing a heavy burden on ER homeostasis (36), making PDAC an ideal context to interrogate this axis. Oncogenic KRAS mutations, present in -90% of PDAC cases, hijack regenerative programs to trigger acinar-to-ductal metaplasia (ADM), yielding pancreatic intraepithelial neoplasia (PanIN) lesions that progress to invasive carcinoma (37–39). Although TRIM21 is reported to promote tumor progression and therapy resistance in established PDAC (21, 40), its role during oncogene-driven initiation remains undefined.

Analysis of Clinical Proteomic Tumor Analysis Consortium (CPTAC) datasets using ProteoCancer Analysis Suite (PCAS) (41) revealed that, unlike most cancers, pancreatic tumors show suppressed BiP and PERK protein despite elevated transcript levels of core UPR genes (BiP (HSPA5), PERK (EIF2AK3), ATF6 and ATF4), and Immunohistochemistry (IHC) on the Human Protein Atlas (https://v24.proteinatlas.org/) independently confirmed reduced BiP with elevated TRIM21 in PDAC (Figure S8A-S8E). Consistent with direct antagonism, TRIM21 and BiP protein levels were negatively correlated in PDAC (Figure S8F). In *Kras*^G12D/+^; *Pdx1*^Cre/+^ (KC) mice, TRIM21 protein progressively increased with tumor advancement, whereas adaptive UPR regulators (ATF6, PERK and IRE1α) and effectors (spliced XBP1, CHOP and EIF2A) declined reciprocally (Figure S8G and S8H).

To establish causality, we crossed KC mice with a pancreas-specific *Trim21* knockout (named *Trim21*^fl/fl^; KC) (Figure 5A and S8I), confirming efficient ablation in pancreatic tissue (Figure 5B and 5C). *Trim21*^fl/fl^; KC mice exhibited delayed tumorigenesis and significantly reduced pancreas-to-body weight ratios at 6 months relative to KC controls (Figure 5D and 5E).

**Figure 5.**
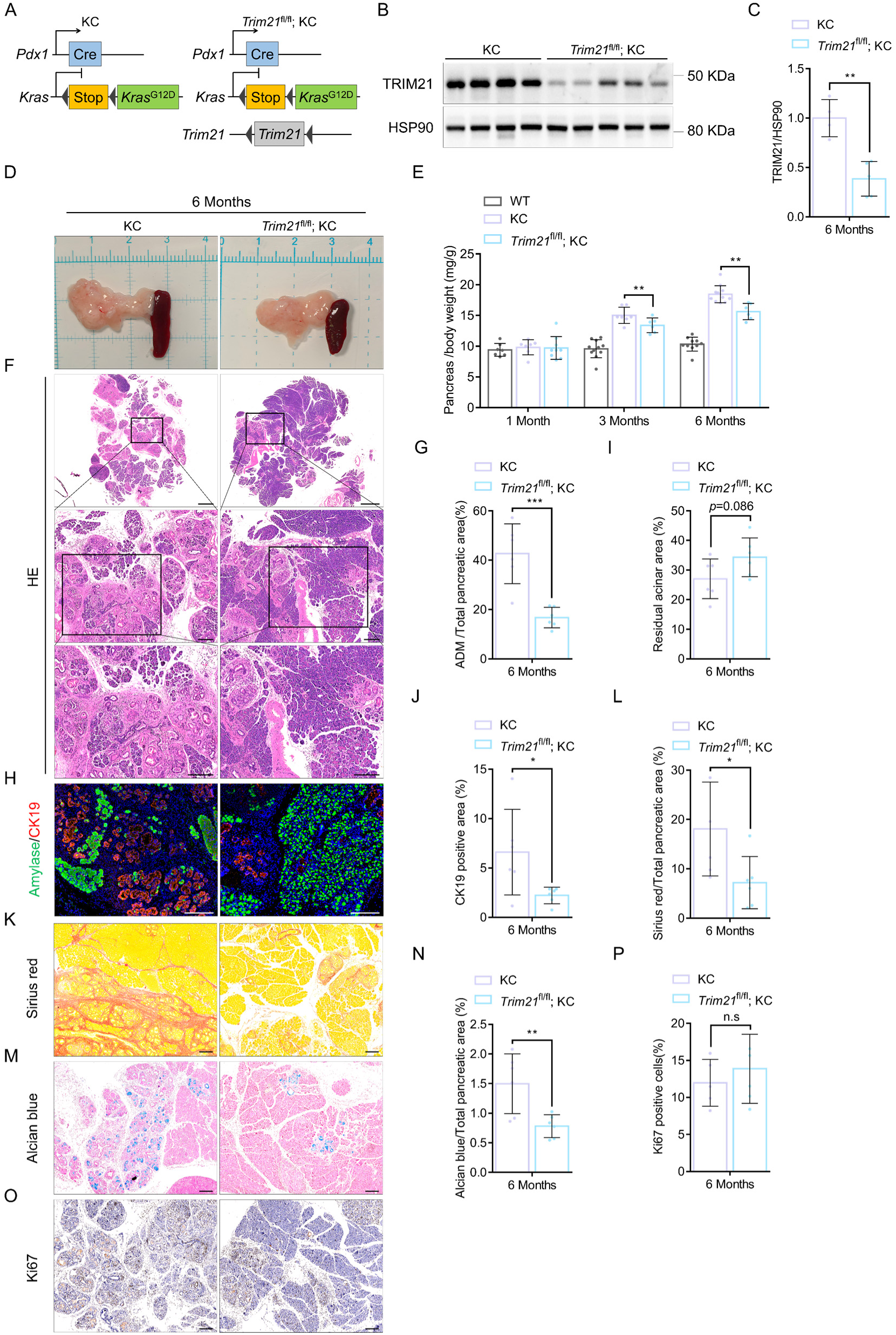
Ablation of *Trim21* in pancreatic cells inhibits PDAC tumorigenesis in KC mice. (A) Generation strategies for the KC and *Trim21*^fl/fl^; KC mouse models. (B and C) TRIM21 protein expression in pancreatic tissues from 6-month-old KC and *Trim21*^fl/fl^; KC mice was assessed by Western blotting, with HSP90 as a loading control (B); blots were subjected to densitometric quantification (C). Data were presented as mean ± SD for 4-5 mice per genotype. **: *P*<0.01 (unpaired Student’s two-tailed t-test). (D) Representative pancreatic tissue morphology in 6-month-old KC and *Trim21*^fl/fl^; KC mice. Scale bar, 1 cm. (E) Pancreas-to-body weight ratio in WT, KC, and *Trim21*^fl/fl^; KC mice at 1, 3, and 6 months of age. Data were presented as mean ± SD for 6-11 mice per genotype. **: *P*<0.01 (one-way ANOVA with Dunnett’s multiple-comparisons test). (F and G) Hematoxylin and eosin (H&E) staining of pancreatic sections from 6-month-old KC and *Trim21*^fl/fl^; KC mice, shown at 2×, 10×, and 20× magnifications. Scale bars, 1,000, 200, and 200 μm, respectively (F). Quantification of ADM as a percentage of total pancreatic area (G). Data were presented as mean ± SD for 6 mice per genotype. ***: *P*<0.001 (unpaired Student’s two-tailed t-test). (H-J) Immunofluorescence analysis of amylase and CK19 in pancreatic sections from 6-month-old KC and *Trim21*^fl/fl^; KC mice (H). Amylase and CK19 served as markers for acinar and ductal cells, respectively. Scale bar, 200 μm. Quantification of amylase (I) and CK19 (J) as a percentage of total pancreatic area. Data were presented as mean ± SD for 6 mice per genotype. *: *P*<0.05 (unpaired Student’s two-tailed t-test). (K-P) Sirius red staining (K), Alcian blue/eosin dual staining (M) and Ki67 immunohistochemistry (IHC) (O) of pancreatic sections from 6-month-old KC and *Trim21*^fl/fl^; KC mice. Scale bar, 200 μm. Quantification of Sirius red-positive collagen area (L), Alcian blue-positive areas (N) and Ki67 positive cells (P), each expressed as a percentage of total pancreatic area. Data were presented as mean ± SD for 5-6 mice per genotype. *: *P*<0.05, **: *P*<0.01. ns, not significant (unpaired Student’s two-tailed t-test).

Histologically, *Trim21*^fl/fl^; KC mice displayed markedly attenuated ADM, collagen fibrosis, and pathological mucin deposition (Figure 5F-5N). Notably, this suppression was independent of proliferation, as Ki67 staining showed no difference between genotypes (Figure 5O and 5P). We propose that oncogenic KRAS normally exploits TRIM21 to safeguard the “proteostatic window” required for acinar reprogramming; TRIM21 loss triggers BiP solidification and a dysregulated UPR that halts ductal morphogenesis at its earliest stage. Collectively, these findings identify TRIM21 as a critical survival factor enabling pancreatic cells to withstand the demands of carcinogenesis.

### 6. Ablation of *Trim21* in pancreatic cells disrupts adaptive UPR in ADM cells

Mechanistic analysis of the TRIM21–BiP axis revealed robust activation of adaptive UPR in pancreatic tissues of 6-month-old *Trim21*^fl/fl^; KC mice relative to KC controls (Figure 6A and 6B). Consistent with our in vitro MEF findings, BiP and TRIM21 markedly colocalized in the ADM of KC mice (Figure 6C). To determine whether TRIM21 modulates BiP’s association with UPR sensors, we examined BiP colocalization with PERK and ATF6 in vivo. While KC mice exhibited robust BiP–PERK and BiP–ATF6 colocalization, *Trim21*^fl/fl^; KC mice showed a pronounced reduction, indicating partial dissociation of BiP from PERK/ATF6-containing complexes (Figure 6D and 6E).

**Figure 6.**
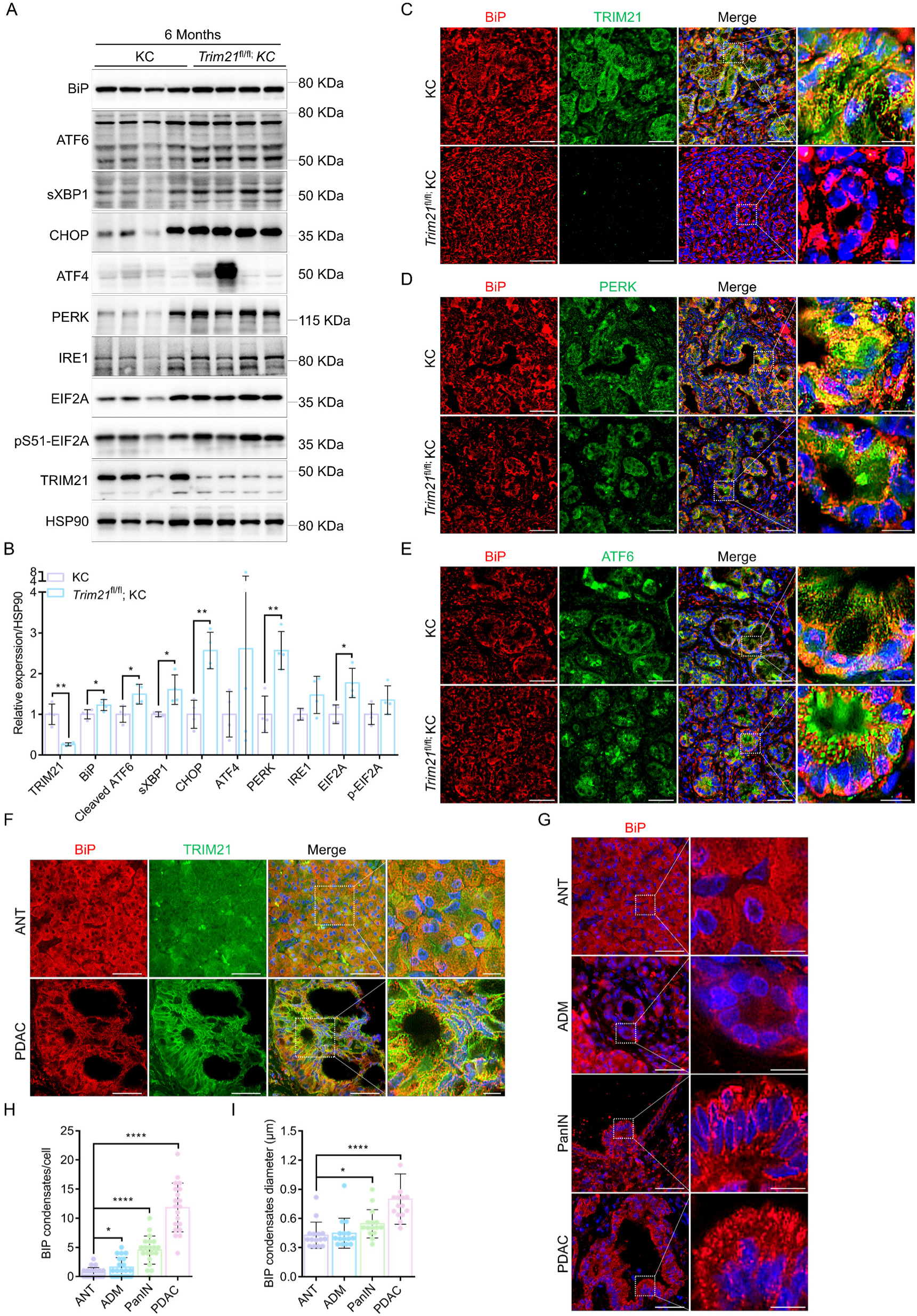
Ablation of *Trim21* in pancreatic cells disrupts adaptive UPR in ADM cells. (A and B) Protein levels of BiP, ATF6, sXBP1, CHOP, ATF4, PERK, IRE1, EIF2A, p-EIF2A, and TRIM21 in the pancreas of 6-month-old KC and *Trim21*^fl/fl^; KC mice were assessed by Western blotting, with HSP90 as a loading control (A); blots were subjected to densitometric quantification (B). Data were presented as mean ± SD for 4 mice per genotype. *: *P*<0.05, **: *P*<0.01 (unpaired Student’s two-tailed t-test). (C-E) Immunofluorescence analysis of BiP together with TRIM21 (C) or PERK (D) or ATF6 (E) in pancreatic sections from 6-month-old KC and *Trim21*^fl/fl^; KC mice. Scale bar, 50 μm and 10 μm. (F) Immunofluorescence analysis of BiP and TRIM21 in pancreatic sections from human adjacent normal tissue (ANT) and PDAC. Scale bar, 50 μm and 10 μm. (G) Immunofluorescence analysis of BiP in pancreatic sections from human ANT, ADM, PanIN and PDAC. Scale bar, 50 μm and 10 μm. (H and I) Quantification of BiP condensate number (H) and diameter (I) for the samples in (G). Data were presented as mean ± SD for 20 cells (H) and 15 cells (G) from 3 independent human PDAC tissues. *: *P*<0.05, ****: *P*<0.0001 (unpaired Student’s two-tailed t-test).

To assess clinical relevance, we analyzed human PDAC tissue microarrays (HPanA030PG03). Compared with adjacent normal tissues (ANT), PDAC samples displayed strong TRIM21–BiP colocalization (Figure 6F). Along the ANT-ADM-PanIN-PDAC progression, BiP condensates increased in both number and diameter (Figure 6G-6I), recapitulating our in vitro and in vivo observations. Collectively, convergent evidence from cellular assays, mouse models, and clinical samples demonstrates that loss of TRIM21 promotes BiP condensate formation, disrupts its interaction with UPR sensors, and perturbs adaptive UPR homeostasis in pancreatic cancer cells.

### 7. *Trim21* inactivation inhibits oncogenic *Kras*-mediated pancreatic cancer by regulating ADM

To interrogate whether the TRIM21–BiP axis directly governs acinar-to-ductal transdifferentiation, we employed a 3D primary acinar organoid model (Figure 7A). *Kras*^G12D^-induced ductal morphogenesis was markedly attenuated in *Trim21*^fl/fl^; KC organoids compared with KC controls (formation efficiency: KC 3.80 ± 0.98% vs *Trim21*^fl/fl^; KC 1.24 ± 1.22% at day 2; KC 13.09 ± 5.06% vs *Trim21*^fl/fl^; KC 5.66 ± 2.04% at day 5; P < 0.05) (Figure 7B and 7C). The ER stress inducer thapsigargin suppressed tubular formation in both genotypes, whereas the inhibitor 4-phenylbutyric acid (4-PBA) rescued the ductal morphogenesis defect in *Trim21*-deficient organoids (Figure 7D and 7E). Consistently, qPCR analysis showed that thapsigargin induced *Trim21*, *Ddit3* and acinar fate factors (*Bhlha15*, *Ptf1a*, *Nr5a2*) while repressing ductal markers (*Hnf6*, *Sox9*, *Fosl1*), effects reversed by 4-PBA (Figure 7F). Furthermore, inhibiting VCP—the engine of ERAD-driven BiP turnover—with NMS-873 strongly reduced duct formation (Figure 7G and 7H).

**Figure 7.**
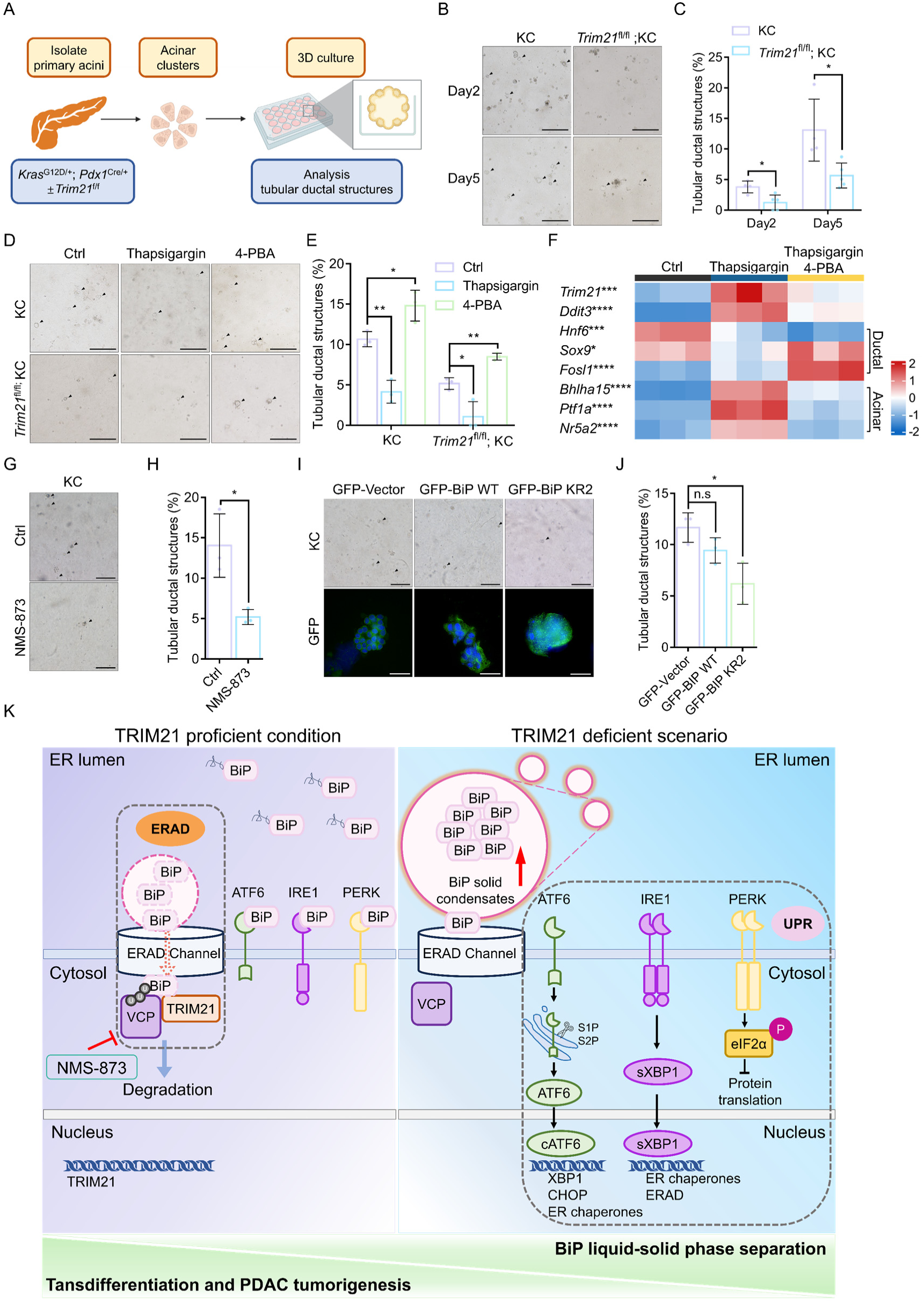
***Trim21* inactivation inhibits oncogenic *Kras*-mediated pancreatic cancer by regulating ADM**. (A) Scheme of the experimental design (created with BioRender). (B) Bright-field images of primary acinar cell clusters from 2-month-old KC and *Trim21*^fl/fl^; KC mice cultured in collagen-based 3D culture at days 2 and 5 after seeding. Arrows indicate ADM cells. Scale bar, 200 μm. (C) Quantification of tubular ductal structures as a percentage of total cells shown in (B). Data were presented as mean ± SD for 4-5 fields of 3 independent experiments. *: *P*<0.05 (unpaired Student’s two-tailed t-test). (D) Bright-field images of primary acinar cell clusters from 2-month-old KC and *Trim21*^fl/fl^; KC mice cultured in collagen-based 3D culture after treatment with DMSO (Ctrl), 0.2 μM Thapsigargin, or 2 mM 4-Phenylbutyric acid (4-PBA) for 48h at day 3 post-seeding. Arrows indicate ADM cells. Scale bar, 200 μm. (E) Quantification of tubular ductal structures as a percentage of total cells shown in (D). Data were presented as mean ± SD of 3 independent experiments. *: *P*<0.05, **: *P*<0.01 (one-way ANOVA with Dunnett’s multiple comparisons test). (F) qPCR analysis of *Trim21*, *Ddit3*, and ductal (*Hnf6*, *Sox9* and *Fosl1*), and acinar (*Bhlha15*, *Ptf1a* and *Nr5a2*) transcription factors in 266-6 cells treated with DMSO (Ctrl), 0.2 μM Thapsigargin, or 0.2 μM Thapsigargin + 2 mM 4-PBA. Expression levels were normalized to *Actb*. *: *P*<0.05, ***: *P*<0.001, ****: *P*<0.0001 (one-way ANOVA with Dunnett’s multiple comparisons test). (G) Bright-field images of primary acinar cell clusters from 2-month-old KC mice cultured in collagen-based 3D culture after treatment with DMSO (Ctrl) or 2 μM NMS-873 for 48h at day 3 post-seeding. Arrows indicate ADM cells. Scale bar, 200 μm. (H) Quantification of tubular ductal structures as a percentage of total cells shown in (G). Data were presented as mean ± SD of 3 independent experiments. *: *P*<0.05 (unpaired Student’s two-tailed t-test). (I) Bright-field and immunofluorescence images of primary acinar cell clusters from 2-month-old KC mice cultured in collagen-based 3D culture and transduced with GFP, BiP-WT-GFP, or BiP-KR2-GFP. Arrows indicate ADM cells. Scale bar, 200 μm and 20 μm. (J) Quantification of tubular ductal structures as a percentage of total cells shown in (I). Data were presented as mean ± SD of 3 independent experiments. *: *P*<0.05, ns, not significant (unpaired Student’s two-tailed t-test). TRIM21-VCP axis regulates BiP phase state and adaptive UPR in pancreatic acinar cells. TRIM21 prevents BiP liquid-to-solid phase separation and promotes VCP-dependent degradation of BiP solid condensates via the ERAD system, thereby maintaining adaptive UPR homeostasis and accelerating the transdifferentiation of acinar cells and PDAC tumorigenesis.

We next asked whether BiP phase behavior, rather than its abundance, dictates ADM. KC clusters were transduced with GFP, liquid-capable BiP-WT-GFP, or solid-prone BiP-KR2-GFP (Figure 7I). Only the BiP-KR2 mutant reduced ductal structures (Figure 7J), indicating that maintaining BiP in its liquid phase (TRIM21-dependent) is required for *Kras*-driven transdifferentiation.

Collectively, oncogenic *Kras* upregulates TRIM21, which ubiquitinates BiP and promotes clearance of solid BiP aggregates via the VCP–ERAD pathway, preserving soluble BiP that engages the ER stress sensors ATF6, PERK and IRE1α to sustain adaptive UPR homeostasis and drive transformation. In the absence of TRIM21, *Kras*- and stress-induced liquid-to-solid phase separation of BiP generates condensates that release these sensors, perturbing adaptive UPR and delaying carcinogenesis. These findings establish the TRIM21–BiP axis as a central regulator of adaptive UPR homeostasis and highlight it as a promising therapeutic target in pancreatic cancer (Figure 7K).

## DISCUSSION

BiP, the primary Hsp70-family chaperone of the ER, is the central coordinator of ER proteostasis and the UPR. Recent in vitro studies have formalized that BiP inherently undergoes liquid-liquid phase separation (LLPS) (16) and that these condensates risk pathological solidification under prolonged stress, such as in the presence of Aβ40 (17). However, a critical conceptual gap remains: under sustained, extreme ER stress—such as during oncogenic reprogramming—how do mammalian cells actively reverse these solidified chaperone condensates? Without an active disassembly mechanism, the ER proteostasis network would inevitably collapse into an irreversible physical state.

Here, we uncover an active, ubiquitin-directed mechanical extraction circuit that uniquely answers this topological and biophysical conundrum. We identify the cytosolic E3 ligase TRIM21 as a remote phase-state regulator of the ER-luminal chaperone BiP. Under sustained ER stress, BiP partitions into solid-like, functionally inert condensates. We demonstrate that TRIM21 engages BiP exclusively at the cytoplasmic face of the ER membrane, catalyzing noncanonical polyubiquitination (K6-, K27-, K33-, and K63-linked) at the SBDα/lid residues K547/553/554/556. Crucially, this specific ubiquitin topology serves as a recruitment tag for the AAA+ ATPase VCP/p97. VCP then provides the essential mechanical force to extract ubiquitinated BiP through the ERAD channel, physically dissolving the solid condensates and routing them for proteasomal degradation. This mechanism solves the spatial topology barrier of how a cytosolic E3 ligase controls a luminal chaperone, redefining ubiquitination not merely as a degradation signal, but as a mechanical recruitment tag to actively dismantle phase-separated condensates.

The physiological necessity of this TRIM21-BiP phase-control axis is profoundly illustrated in pancreatic carcinogenesis. Pancreatic acinar cells impose an exceptional secretory demand on ER homeostasis. Because BiP is an absolute prerequisite for ADM (7), oncogenic KRAS must preserve a highly functional BiP pool to survive the extreme proteotoxic stress of transdifferentiation. Our data suggest that KRAS hijacks TRIM21 to act as a “biophysical shield.” By constantly clearing solidified BiP via the VCP-ERAD axis, TRIM21 preserves a soluble, liquid-like chaperone pool capable of properly sequestering the UPR sensors ATF6, PERK, and IRE1α. Conversely, upon pancreas-specific *Trim21* ablation, BiP undergoes unchecked liquid-to-solid phase separation. These inert solid condensates fail to sequester the UPR sensors, unleashing an exaggerated, unbalanced UPR that ultimately halts ductal morphogenesis and tumorigenesis (42–47). This phenotype is corroborated by human PDAC tissue microarrays, which exhibit a progressive increase in solid BiP condensates alongside TRIM21 up-regulation during malignant transformation.

Importantly, our findings unveil a new pharmacological vulnerability. While targeting transcription or kinase signaling has been the cornerstone of targeted therapy, our study suggests that the physical phase state of organelle chaperones can be therapeutically exploited. By using the VCP inhibitor NMS-873, we successfully dismantled the mechanical extraction axis, artificially forcing BiP into a lethal solid phase and terminating *Kras*-driven ADM in 3D organoids. Thus, the TRIM21-BiP phase condensation axis not only establishes a fundamental paradigm linking ubiquitin-driven proteostasis to biophysical organelle remodeling, but also offers a potent therapeutic window to selectively trigger terminal UPR in early-stage pancreatic cancer.

### Limitations of the study

Several caveats warrant consideration. First, our mechanistic inferences rely primarily on the PANC-1 cell line, MEF cell line and the KC mouse model, which reproduce key genetic lesions but not the full heterogeneity, stromal crosstalk, and immune microenvironment of human tumors. Second, the liquid– solid transition of BiP and its dynamic interaction with UPR sensors remain technically challenging to resolve, and the factors governing this transition are still incompletely defined. Third, although pharmacological ER-stress modulation influences KRAS-driven ADM, the underlying molecular mechanisms remain unclear; reliance on a nonspecific small molecule, without comprehensive dose–response, pharmacokinetic, and pharmacodynamic analyses, limits translational interpretation, and in vivo therapeutic validation is likewise lacking. Future work should prioritize a highly selective BiP phase-state modulator, which would enable dose-titration studies and long-term mechanistic interrogation in genetically engineered models. Such tools will be pivotal in establishing the TRIM21–BiP axis as an actionable target in pancreatic cancer.

## RESOURCE AVAILABILITY

### Lead contact

Further information and requests for resources and reagents should be directed to and will be fulfilled by the lead contact, Wenjie Ge.

### Materials availability

All materials generated in this study are available from the lead contact with a completed materials transfer agreement.

## Data availability

All relevant data to evaluate the conclusions in the paper are within the paper and/or the Supplementary Materials.

## ACKNOWLEDGEMENTS

This work was supported by the National Natural Science Foundation of China (Grant No. 82573779, 82203311), the Jiangsu Province Capability Improvement Project through Science, Technology and Education (NO. CXZX202234), the Open Project of Key Laboratory of Science and Engineering for the Multi-modal Prevention and Control of Major Chronic Diseases, Ministry of Industry and Information Technology (Grant No. MCD-2023-1-01), the Major Project of Basic Science (Natural Science) Research in Colleges and Universities of Jiangsu Province (25KJA180006), the Research Foundation of Xuzhou Medical University (D2023004, D2023012, D2022034, D2025006), “Double Innovation” talents projects of Jiangsu Province, the Basic Science (Natural Science) Research Project of Higher Education Institutions in Jiangsu Province (24KJD320006, 25KJB180016). Jiangsu Postgraduate Research and Practice Innovation Program (KYCX24_3079 and KYCX25_3315). The funders had no role in study design, data collection and analysis, decision to publish, or preparation of the manuscript.

## AUTHOR CONTRIBUTIONS

Conceptualization was the responsibility of W.J.G., Y.F.Z., J.B., Y.F.H. and T.T.W.. Methodology was the responsibility of Y.F.Z., R.H.Y., X.L., Y.H., Q.D.L., Y.R.Z., H.X.Z., X.H.H., Y.X.C., X.Y.W., Z.J.Z., J.K.D., J.X., C.H., L.Y.M., J.Y.W., N.W., J.T.X. and W.H.Z.. The investigation was conducted by W.J.G., Y.F.Z., R.H.Y., X.L. and Y.H.. The original draft was written by Y.F.Z. and R.H.Y.. The reviewing and editing were performed by W.J.G., J.B., Y.F.H., T.T.W. and C.W.. Fund acquisition was carried out by WJ.G, Y.F.Z., X.L. and Y.H.. Supervision was the responsibility of WJ.G., J.B., Y.F.H. and T.T.W.. All authors read and approved the final version of the manuscript.

## DECLARATION OF INTERESTS

All authors declare no competing interests.

---

**Figure S1.**
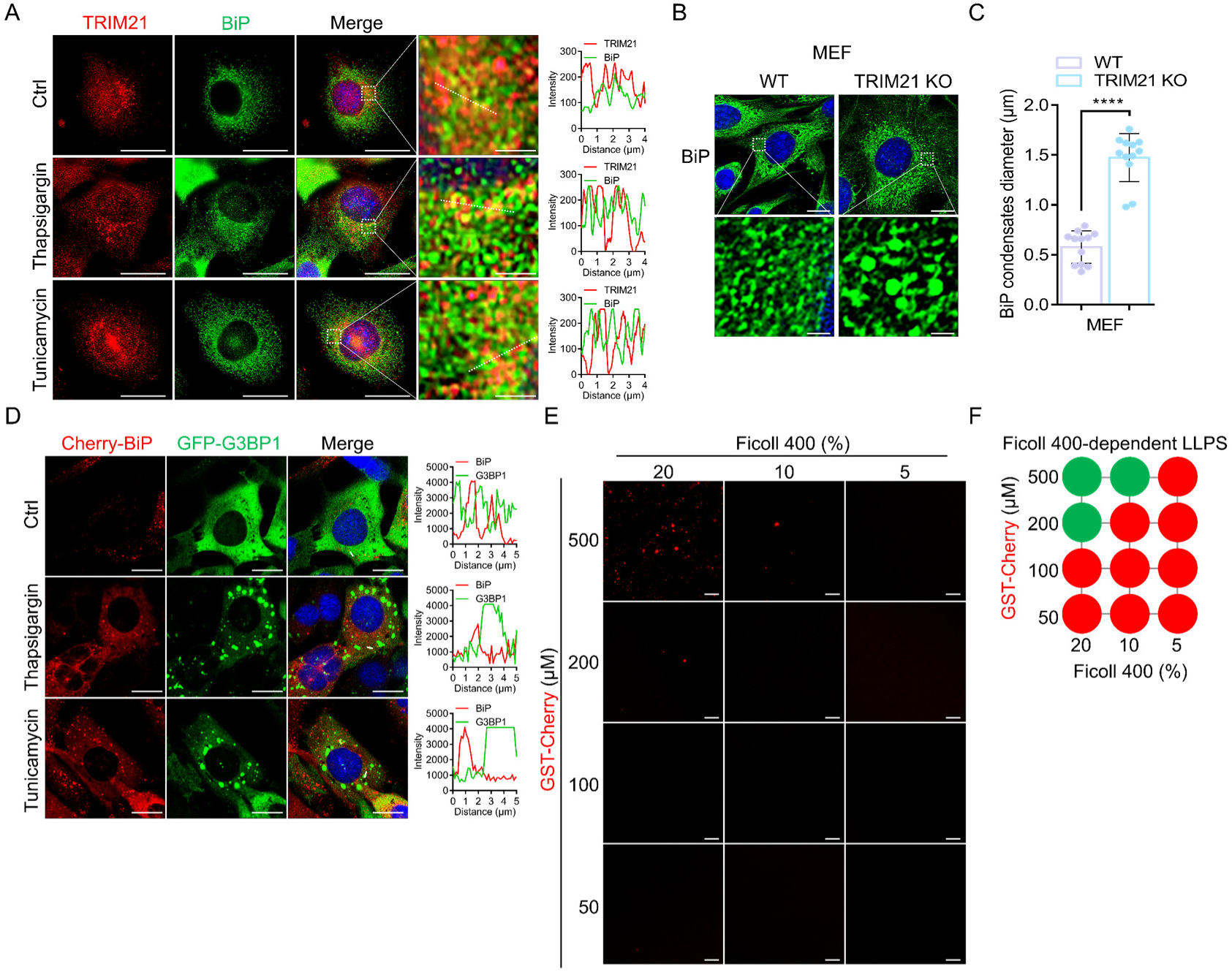
BiP condensates form independently of stress granules. (A) Immunofluorescence analysis of BiP and TRIM21 in MEF cells treated with DMSO (Ctrl), 1 μM Thapsigargin, or 5 μg/mL Tunicamycin for 4 hours. Scale bar, 20 μm and 2 μm. (B and C) Immunofluorescence analysis of BiP in WT MEF and TRIM21 KO MEF cells treated with 1 μM Thapsigargin for 4 hours (B). Scale bar, 20 μm and 2 μm. Quantification of the diameter of BiP condensates shown in (B); data were presented as mean ± SD for 12 cells from 3 independent experiments. ****: *P*<0.0001 (unpaired Student’s two-tailed t-test). (D) Immunofluorescence analysis of BiP and G3BP1 in WT MEF cells stably expressing Cherry-BiP and GFP-G3BP1, after treatment with DMSO (Ctrl), 1 μM Thapsigargin, or 5 μg/mL Tunicamycin for 4 hours. Scale bar, 20 μm. (E and F) Phase-separated droplets of GST-Cherry imaged in a Ficoll-400-based crowding environment, with concentration-dependent effects documented through representative microscopy (E) and summary of phase separation behaviors of GST-Cherry at the indicated concentrations (F). Scale bar, 20 µm.

**Figure S2.**
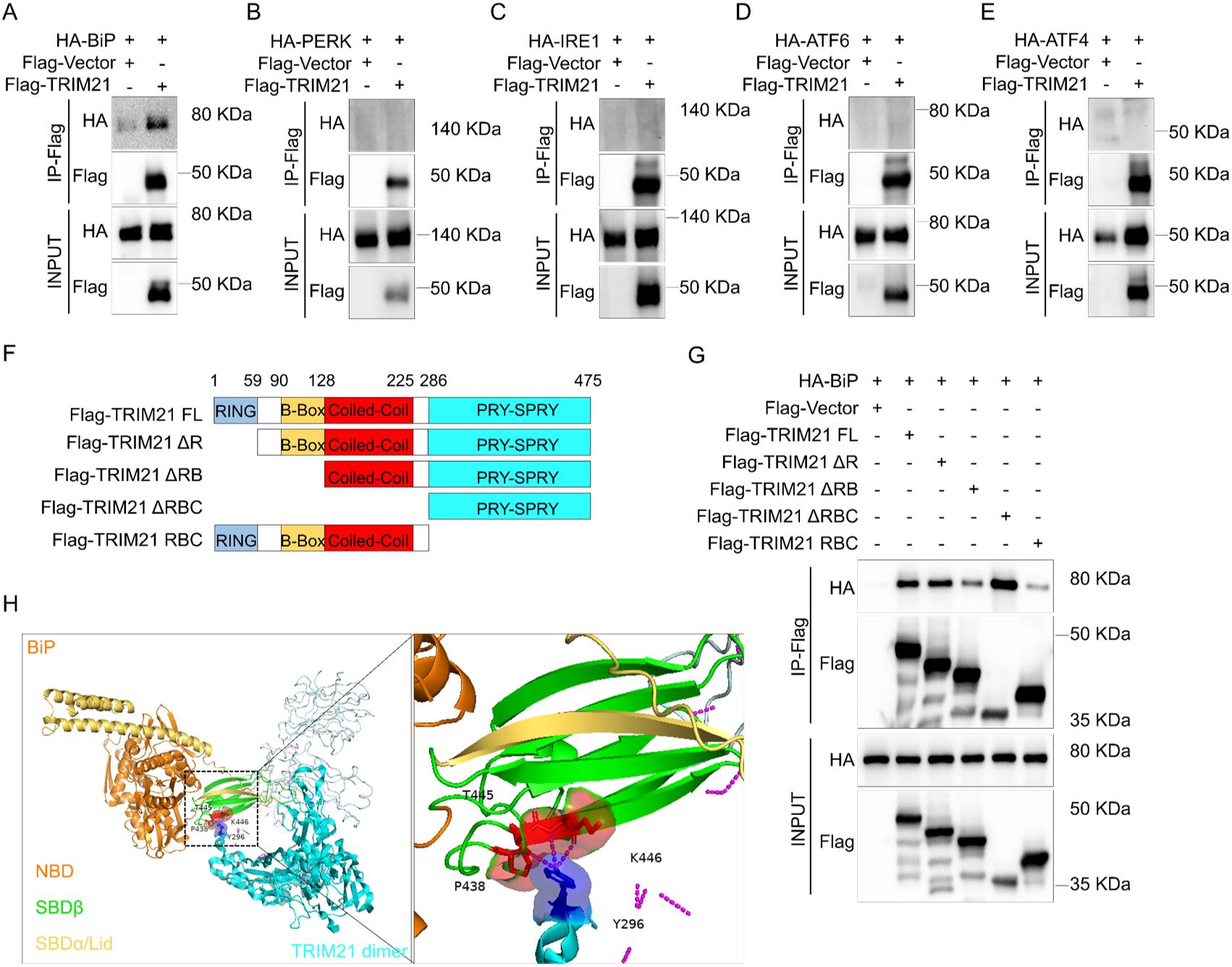
TRIM21 binds directly to BiP via PRY-SPRY domain. (A-E) Co-IP assays in HEK-293T cells co-transfected with HA-BiP (A), HA-PERK (B), HA-IRE1 (C), HA-ATF6 (D), HA-ATF4 (E), together with the indicated combinations of empty Flag-vector and Flag-TRIM21. Immunoprecipitates (top) and inputs (bottom) were analyzed by Western blotting against anti-HA or anti-Flag. (F) Schematic of the cloning strategy used to generate Flag-TRIM21 truncation mutants. (G)Co-IP assays in HEK-293T cells co-transfected with HA-BiP together with full-length TRIM21 or the indicated truncated forms of TRIM21. Immunoprecipitates (top) and inputs (bottom) were analyzed by Western blotting against anti-HA or anti-Flag. (H) Predicted complex structure of the TRIM21 dimer (PDB: 2IWG, cyan) and BiP (PDB: 6ASY, yellow), generated by docking with HDOCK(48). The PRY-SPRY domain of TRIM21 (aa 286-475) and the substrate-binding domain (SBD) of BiP (aa 420-552) are indicated by black dashed boxes. Residues predicted to mediate the interaction are shown as blue (TRIM21) and red (BiP) spheres, with side chains displayed as sticks; predicted hydrogen bonds are shown as magenta dashed lines. The right panel is an enlarged view of the three interactions involving P438, T445, and K446 of BiP with Y296 of TRIM21.

**Figure S3.**
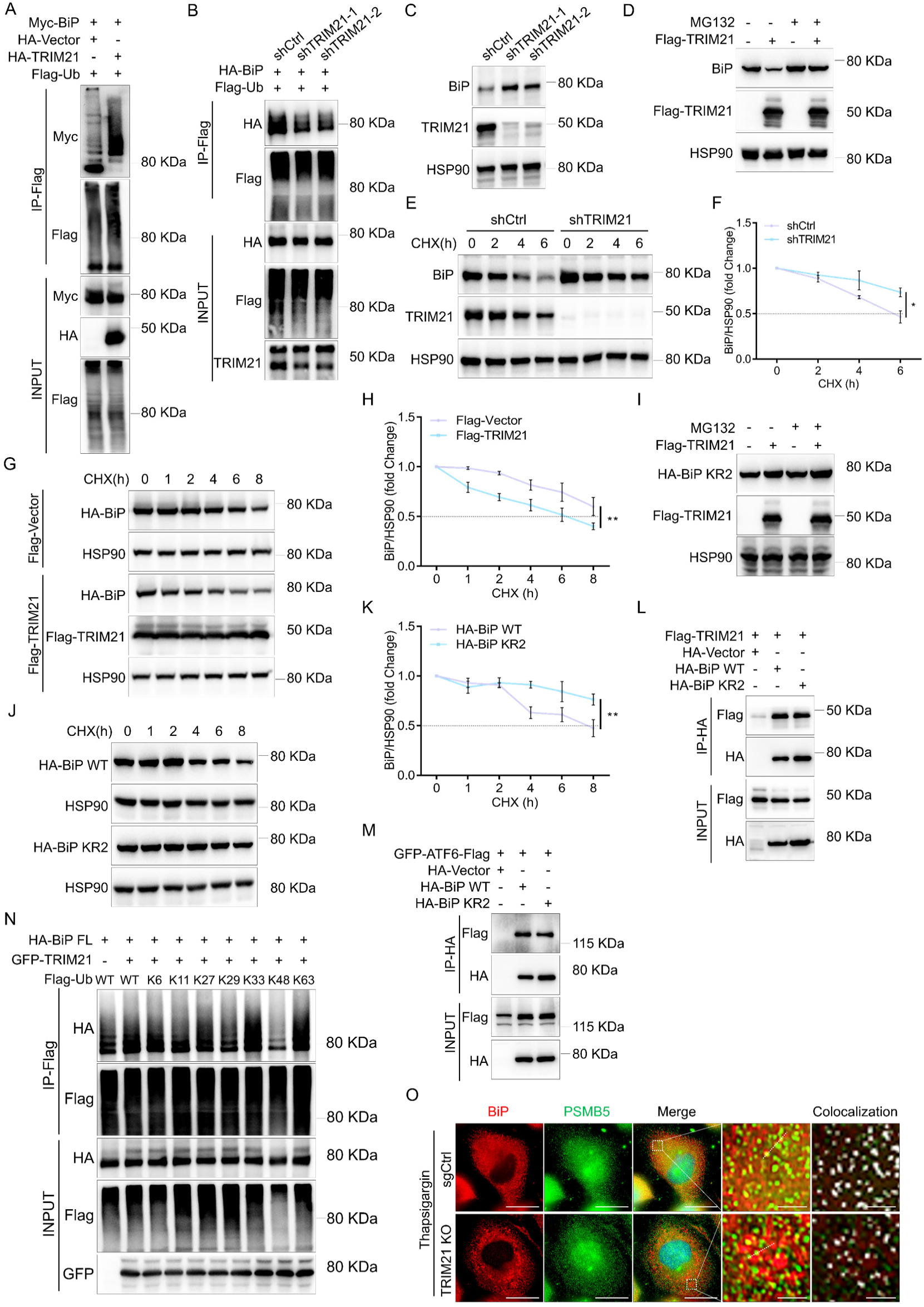
TRIM21 promotes the ubiquitin-mediated degradation of BiP. (A) Ubiquitination assays in HEK-293T cells co-transfected with Myc-BiP and Flag-Ub, along with HA-TRIM21. Samples recovered on Flag-beads (top) and inputs (bottom) were analyzed by Western blotting against anti-Myc, anti-Flag, and anti-HA as indicated. (B) Ubiquitination assays in HEK-293T shCtrl, shTRIM21-1, and shTRIM21-2 cells. Samples recovered on Flag-beads (top) and inputs (bottom) were analyzed by blotting against anti-HA, anti-Flag, and anti-TRIM21 as indicated. (C) Immunoblot analysis of BiP and TRIM21 levels in PANC-1 cells stably expressing shCtrl or TRIM21-targeting shRNAs. HSP90 served as a loading control. (D) Immunoblot analysis of BiP levels in HEK-293T cells transfected with empty Flag vector or Flag-TRIM21, with or without 10 μM MG132 for 6 hours. HSP90 served as a loading control. (E and F) Immunoblot analysis of BiP levels in sgCtrl and TRIM21 KO PANC-1 cells treated with 100 μg/mL CHX to block de novo protein synthesis (E) . Densitometric quantification of BiP levels (F). Data were presented as mean ± SD of 3 independent experiments. *: *P*<0.05 (two-way ANOVA with Tukey’s multiple-comparisons test). (G and H) Immunoblot analysis of BiP levels in HEK-293T cells co-transfected with HA-BiP and Flag-TRIM21, or HA-BiP alone, and treated with 100 μg/mL CHX (G). Densitometric quantification of BiP levels (H). Data were presented as mean ± SD of 3 independent experiments. **: *P*<0.01 (two-way ANOVA with Tukey’s multiple-comparisons test). (I) Immunoblot analysis of BiP KR2 mutant levels in HEK-293T cells transfected with empty Flag vector or Flag-TRIM21, with or without 10 μM MG132 for 6 hours. HSP90 served as a loading control. (J and K) Immunoblot analysis of full-length HA-BiP or the KR2 mutant levels in HEK-293T cells treated with 100 μg/mL CHX (J). Densitometric quantification of BiP levels (K). Data were presented as mean ± SD of 3 independent experiments. **: *P*<0.01 (two-way ANOVA with Tukey’s multiple-comparisons test). (L and M) Co-IP assays in HEK-293T cells co-transfected with Flag-TRIM21 (L) or GFP-ATF6-Flag (M) and the indicated combinations of empty HA-vector, HA-BiP WT, and HA-BiP KR2 mutant. Immunoprecipitates (top) and inputs (bottom) were analyzed by Western blotting against anti-Flag or anti-HA. (N) Ubiquitination assay in HEK-293T cells co-transfected with Flag-Ub WT or the indicated Flag-Ub mutants, along with HA-BiP and GFP-TRIM21. Samples recovered on Flag-beads (top) and inputs (bottom) were analyzed by Western blotting against anti-HA, anti-Flag, and anti-GFP as indicated. (O)Immunofluorescence analysis of BiP and PSMB5 in sgCtlrl and TRIM21 KO PANC-1 cells treated with 1 μM Thapsigargin for 4 hours. Colocalization was analyzed by ImageJ. Scale bar, 20 μm and 2 μm.

**Figure S4.**
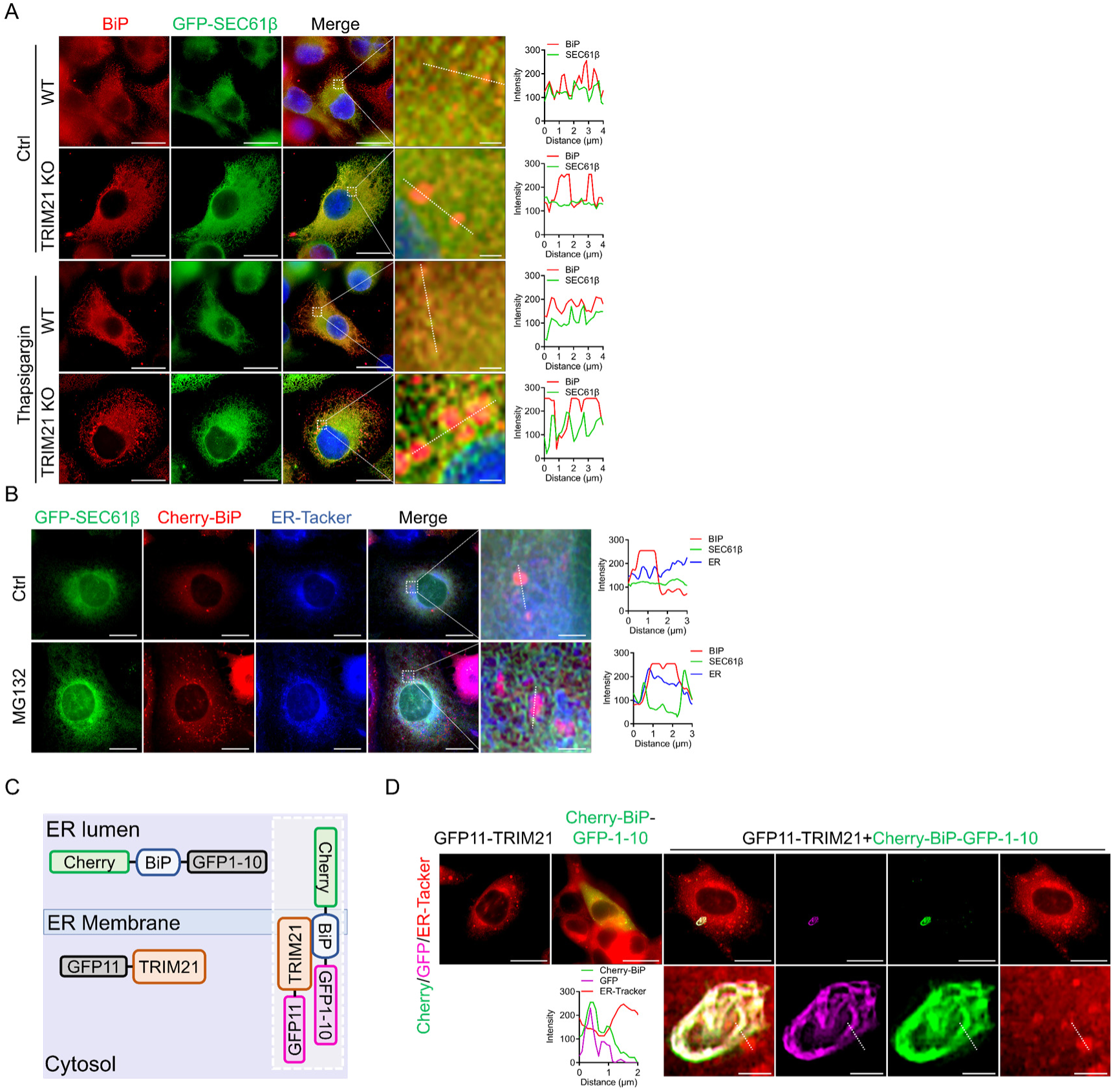
TRIM21 engages BiP on the cytosolic face of the ER membrane. (A) Immunofluorescence analysis of endogenous BiP and GFP-SEC61β in WT and TRIM21 KO MEF cells stably expressing GFP-SEC61β after treated with DMSO (Ctrl) or 1 μM Thapsigargin for 4 hours. Scale bar, 20 μm and 2 μm. (B) Immunofluorescence analysis of ER-Tacker, Cherry-BiP and GFP-SEC61β in WT MEF cells treated with 10 μM MG132 for 6 hours. Scale bar, 20 μm and 2 μm. (C and D) Schematic illustration of the split-GFP (GFP1-10/GFP11) bimolecular fluorescence complementation (BiFC) system used to visualize TRIM21-BiP proximity (C). Cherry-BiP-GFP1-10 and GFP11-TRIM21, when expressed separately, do not emit green fluorescence. When ER luminal Cherry-BiP-GFP1-10 and cytosolic GFP11-TRIM21 were brought into spatial proximity, the split GFP fragments reconstitute an intact fluorophore and generate a green fluorescence signal, reporting the interaction between BiP and TRIM21 on the cytosolic face of the ER (D). Scale bar, 20 μm and 2 μm.

**Figure S5.**
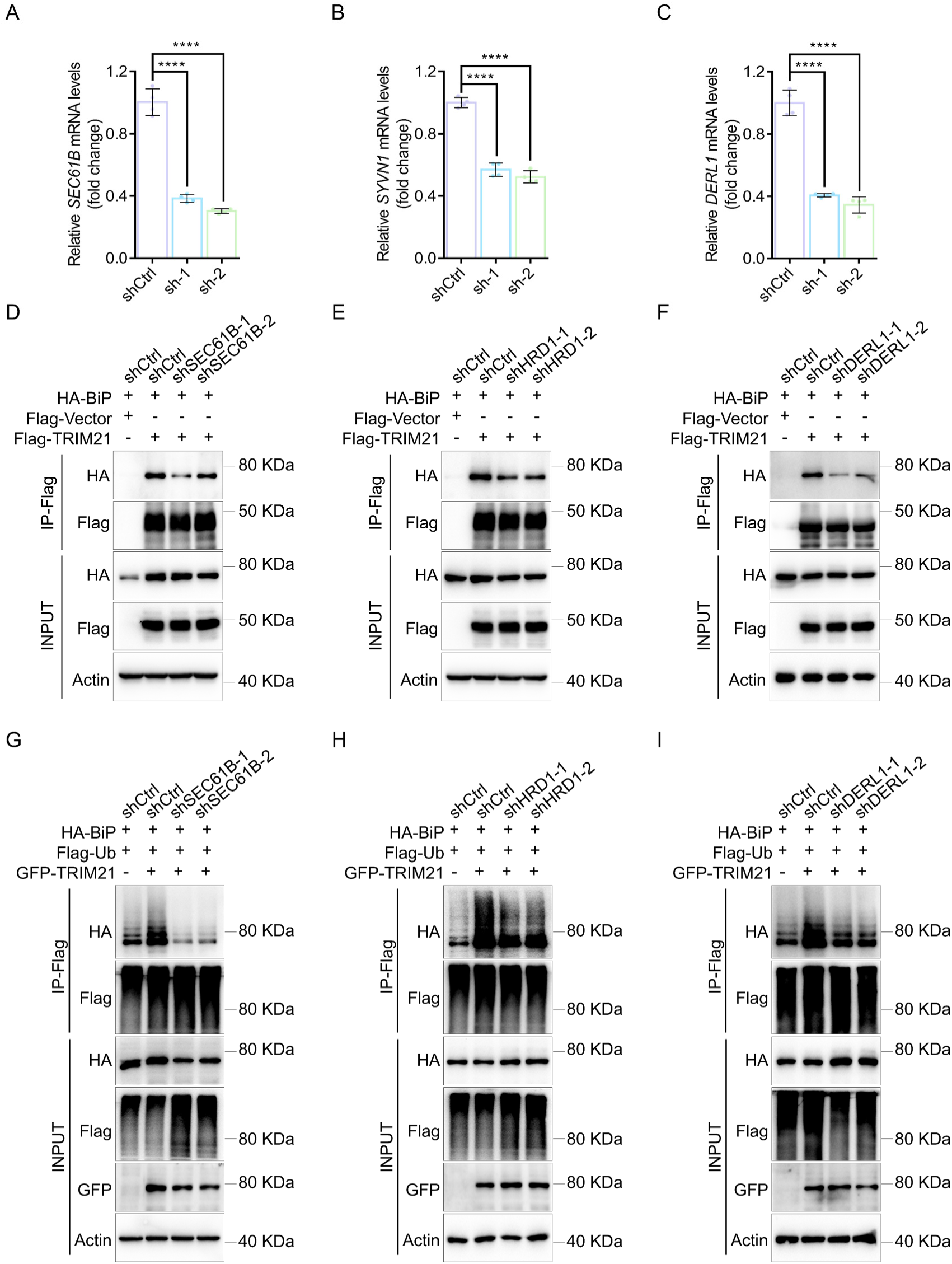
TRIM21 promotes BiP ubiquitination in an ERAD-dependent manner. (A-C) qPCR analysis of knockdown efficiency in HEK293T cells stably expressing the indicated shRNAs targeting *SEC61B* (A), *SYVN1* (B), and *DERL1* (C). Expression levels were normalized to *Actb*. Data were presented as mean ± SD of 4 independent experiments.****: *P*<0.0001 (unpaired Student’s two-tailed t-test). (D-F) Co-IP assays in HEK-293T shCtrl, shSEC61B (D), shHRD1/*SYVN1* (E) or shDERL1 (F) cells co-transfected with HA-BiP and Flag-TRIM21. Immunoprecipitates (top) and inputs (bottom) were analyzed by Western blotting against anti-HA, anti-Flag, and anti-Actin. (G-I) Ubiquitination assays in HEK-293T shCtrl, shSEC61B (G), shHRD1 (H) or shDERL1 (I) cells. Samples recovered on Flag-beads (top) and inputs (bottom) were analyzed by blotting against anti-HA, anti-Flag, anti-GFP, and anti-Actin as indicated.

**Figure S6.**
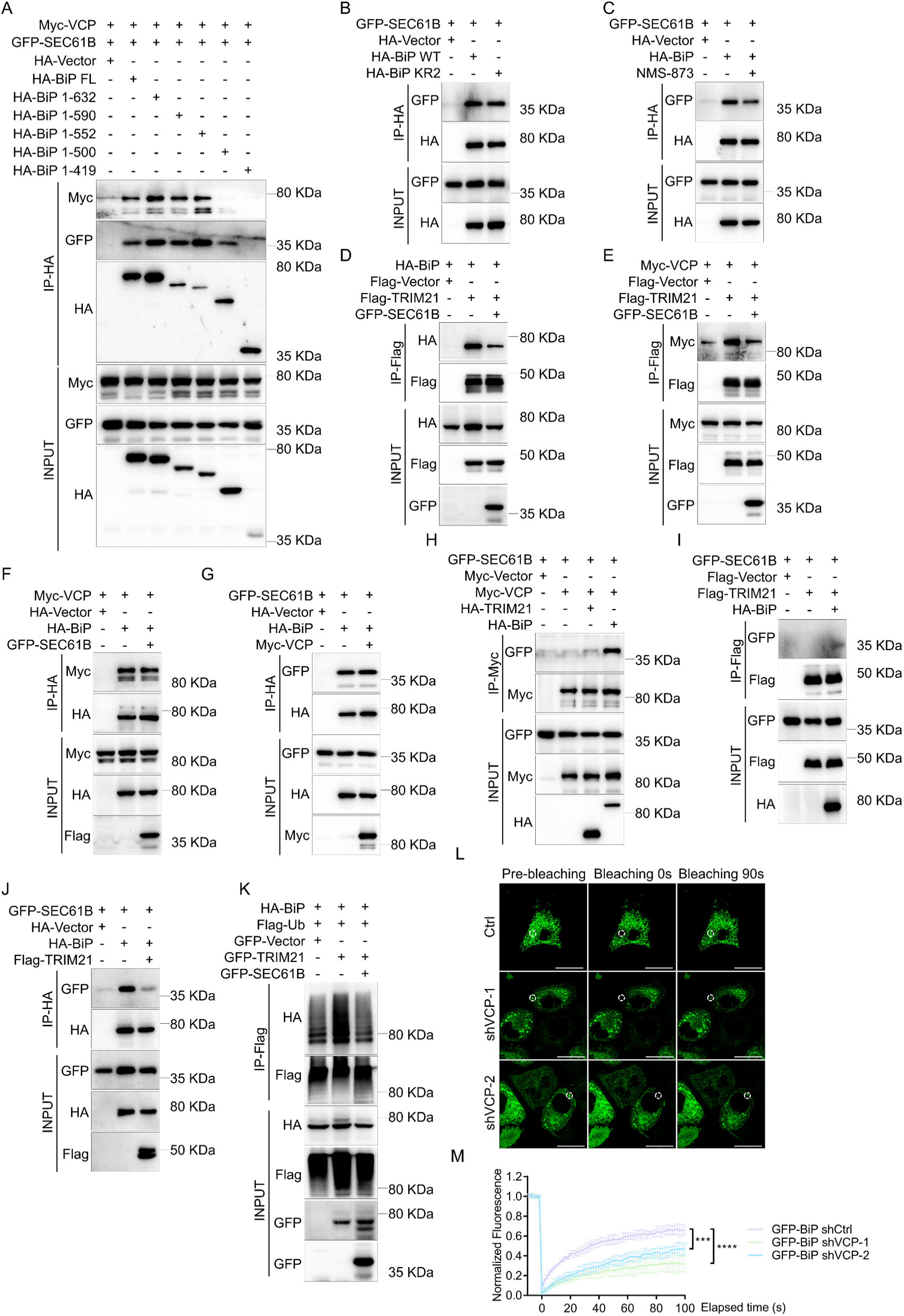
TRIM21 and VCP compete with SEC61β for binding to BiP. (A) Co-IP assays in HEK-293T cells co-transfected with Myc-VCP and GFP-SEC61B and either full-length BiP or the indicated BiP truncation mutants. Immunoprecipitates (top) and inputs (bottom) were analyzed by Western blotting against anti-Myc, anti-GFP or anti-HA. (B) Co-IP assays in HEK-293T cells co-transfected with GFP-SEC61B and the indicated combinations of empty HA-vector, HA-BiP WT, and HA-BiP KR2 mutant. Immunoprecipitates (top) and inputs (bottom) were analyzed by Western blotting against anti-GFP or anti-HA. (C) Co-IP assays in HEK-293T cells co-transfected with GFP-SEC61B and HA-BiP, treated with DMSO (Ctrl) or 2 μM NMS-873 for 4 hours. Immunoprecipitates (top) and inputs (bottom) were analyzed by Western blotting against anti-HA and anti-GFP. (D) Co-IP assays in HEK-293T cells co-transfected with HA-BiP and Flag-TRIM21, with or without GFP-SEC61B. Immunoprecipitates (top) and inputs (bottom) were analyzed by Western blotting against anti-HA, anti-Flag, and anti-GFP. (E) Co-IP assays in HEK-293T cells co-transfected with Myc-VCP and Flag-TRIM21, with or without GFP-SEC61B. Immunoprecipitates (top) and inputs (bottom) were analyzed by Western blotting against anti-Myc, anti-Flag, and anti-GFP. (F) Co-IP assays in HEK-293T cells co-transfected with Myc-VCP and HA-BiP, with or without GFP-SEC61B. Immunoprecipitates (top) and inputs (bottom) were analyzed by Western blotting against anti-Myc, anti-HA, and anti-GFP. (G) Co-IP assays in HEK-293T cells co-transfected with GFP-SEC61B and HA-BiP, with or without Myc-VCP. Immunoprecipitates (top) and inputs (bottom) were analyzed by Western blotting against anti-GFP, anti-HA, and anti-Myc. (H) Co-IP assays in HEK-293T cells co-transfected with GFP-SEC61B and Myc-VCP, with or without HA-TRIM21 or HA-BiP. Immunoprecipitates (top) and inputs (bottom) were analyzed by Western blotting against anti-GFP, anti-Myc, and anti-HA. (I) Co-IP assays in HEK-293T cells co-transfected with GFP-SEC61B and Flag-TRIM21, with or without HA-BiP. Immunoprecipitates (top) and inputs (bottom) were analyzed by Western blotting against anti-GFP, anti-Flag, and anti-HA. (J) Co-IP assays in HEK-293T cells co-transfected with GFP-SEC61B and HA-BiP, with or without Flag-TRIM21. Immunoprecipitates (top) and inputs (bottom) were analyzed by Western blotting against anti-GFP, anti-HA, and anti-Flag. (K) Ubiquitination assay in HEK-293T cells co-transfected with Flag-Ub WT, HA-BiP and GFP-TRIM21, with or without GFP-SEC61B. Samples recovered on Flag-beads (top) and inputs (bottom) were analyzed by blotting against anti-HA, anti-Flag, and anti-GFP. (L) PANC-1 shCtrl, shVCP-1 and shVCP-2 cells transduced with lenti-EFs-BiP-GFP were subjected to FRAP analysis on a Leica STELLARIS 5 confocal microscope (488 nm channel). Time-lapse imaging captured BiP-GFP condensate recovery. Scale bar, 20 μm. (M) Quantification of FRAP recovery of BiP-GFP condensates in PANC-1 shCtrl, shVCP-1 and shVCP-2 cells, presented as mean ± SD of 4 bleached areas. ***: *P*<0.001, ****: *P*<0.0001 (two-way ANOVA with Tukey’s multiple-comparisons test).

**Figure S7.**
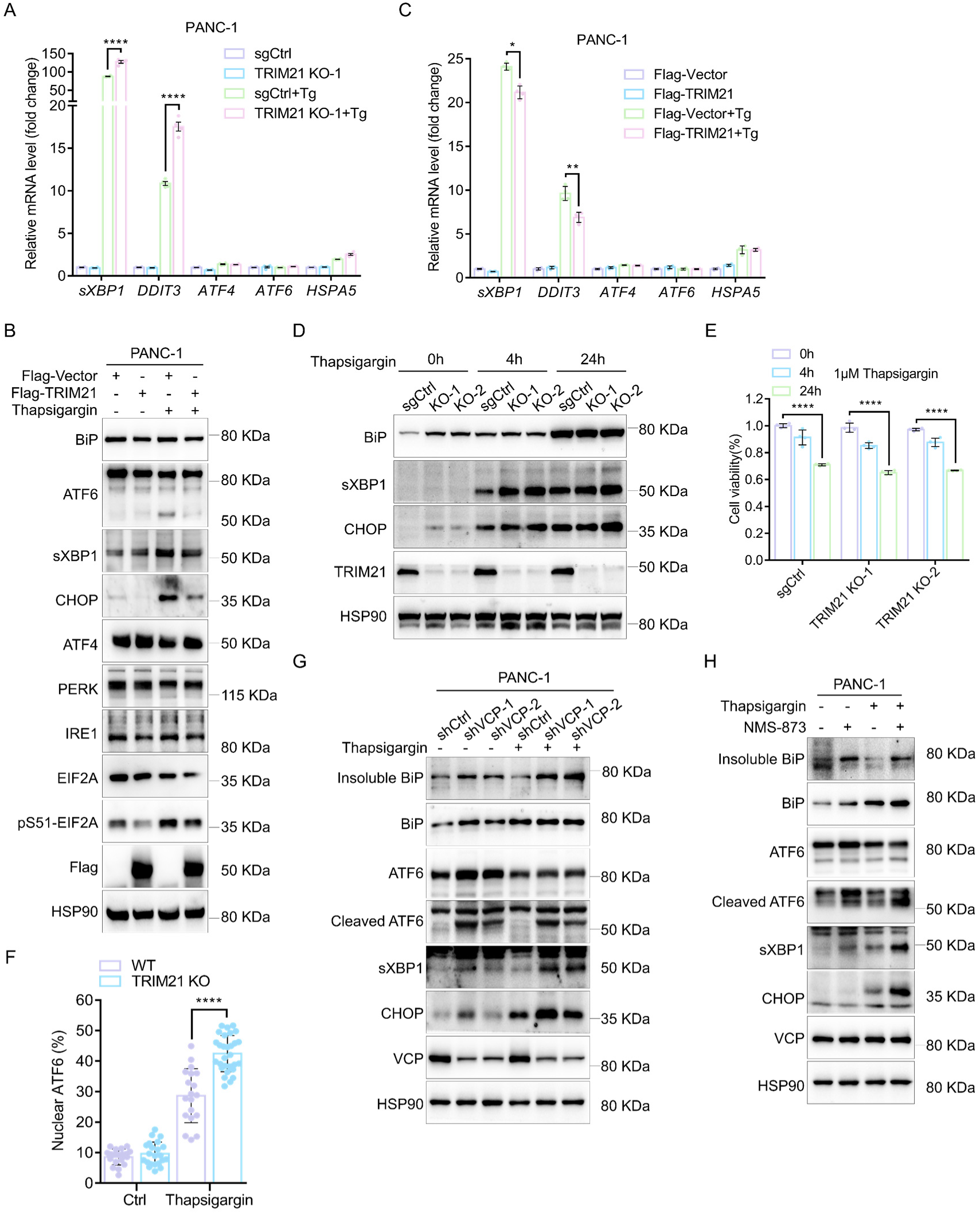
TRIM21 and VCP act as an adaptive UPR regulator in PDAC. (A) qPCR analysis of relative s*XBP1*, *DDIT3*, *ATF4*, *ATF6* and *HSPA5* mRNA levels in PANC-1 sgCtrl and TRIM21 KO cells treated with DMSO (Ctrl) or 1 μM Thapsigargin for 4 hours. Expression levels were normalized to *Actb*. Data were presented as mean ± SD of 4 independent experiments. *: *P*<0.05, **: *P*<0.01, ****: *P*<0.0001 (unpaired Student’s two-tailed t-test). (B) Protein levels of BiP, ATF6, sXBP1, CHOP, ATF4, PERK, IRE1, EIF2A, pS51-EIF2A, and Flag-TRIM21 in PANC-1 transfected with Flag vector or Flag-TRIM21 cells treated with DMSO (Ctrl) or 1 μM Thapsigargin for 4 hours. HSP90 was used as a loading control. (C) qPCR analysis of relative s*XBP1*, *DDIT3*, *ATF4*, *ATF6* and *HSPA5* mRNA levels in PANC-1 cells transfected with Flag vector or Flag-TRIM21 and treated with DMSO (Ctrl) or 1 μM Thapsigargin for 4 hours. Expression levels were normalized to *Actb*. Data were presented as mean ± SD of 4 independent experiments. *: *P*<0.05, **: *P*<0.01, ****: *P*<0.0001 (unpaired Student’s two-tailed t-test). (D) Protein levels of BiP, sXBP1, CHOP and TRIM21 in PANC-1 sgCtrl and TRIM21 KO cells treated with DMSO (Ctrl) or 1 μM Thapsigargin for 4 or 24 hours. HSP90 was used as a loading control. (E) Proliferation of PANC-1 sgCtrl and TRIM21 KO cells treated with DMSO (Ctrl) or 1 μM Thapsigargin for 4 or 24 hours. ****: *P*<0.0001 (one-way ANOVA with Dunnett’s multiple-comparisons test). (F) Quantification of nuclear ATF6 fluorescence intensity as a percentage of total ATF6 fluorescence intensity in Figure 4C. Data were mean ± SD for 19-29 cells from 3 independent experiments. ****: *P*<0.0001 (unpaired Student’s two-tailed t-test). (G) Protein levels of insoluble BiP, BiP, ATF6, cleaved ATF6, sXBP1, CHOP, and VCP in PANC-1 shCtrl, shVCP-1, and shVCP-2 cells treated with DMSO (Ctrl) or 1 μM Thapsigargin for 4 hours. HSP90 was used as a loading control. (H) Protein levels of insoluble BiP, BiP, ATF6, cleaved ATF6, sXBP1, CHOP, and VCP in PANC-1 cells treated with DMSO (Ctrl), 1 μM Thapsigargin or 2 μM NMS-873 for 4 hours. HSP90 was used as a loading control.

**Figure S8.**
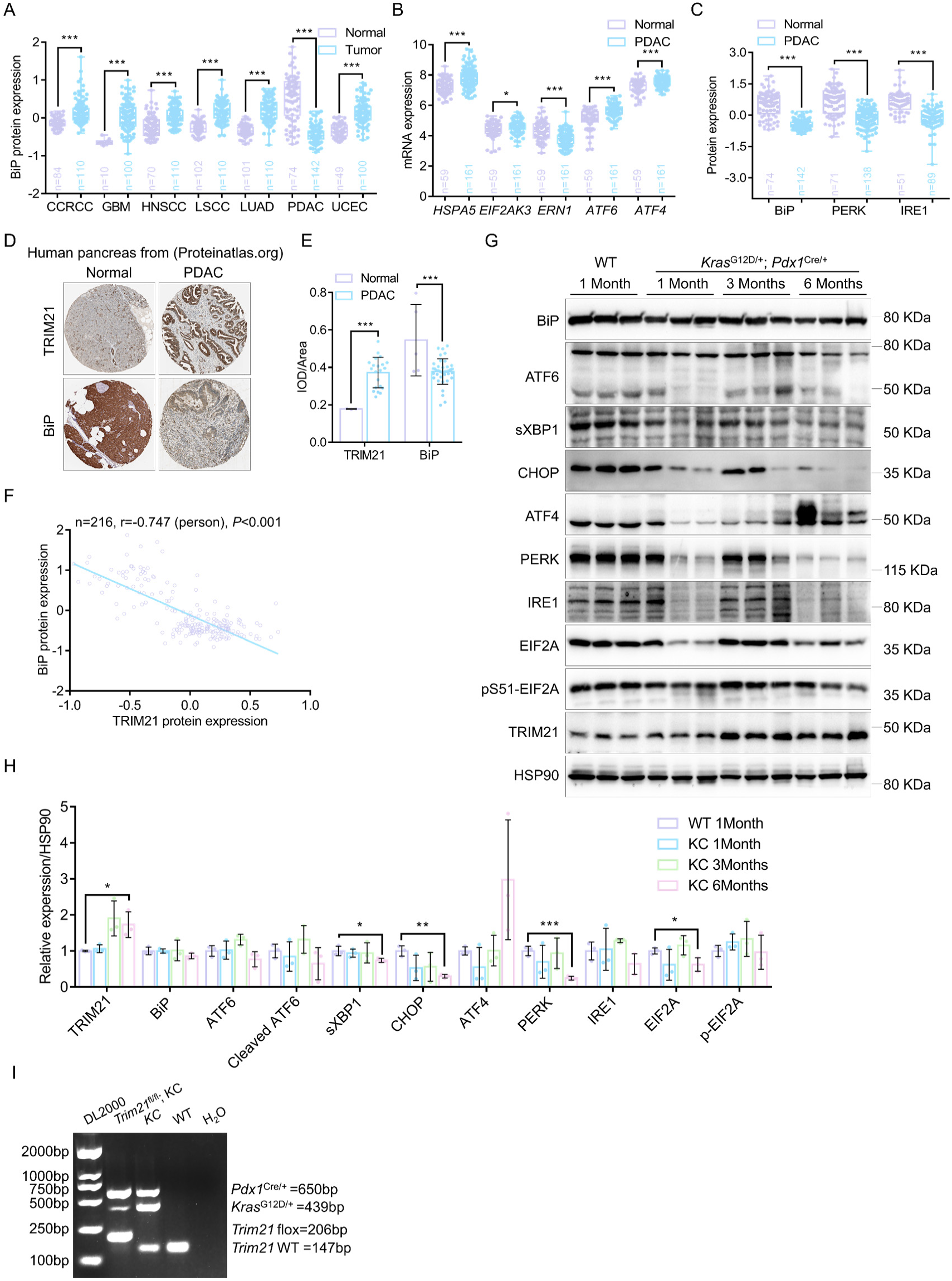
Pancreatic tumorigenesis in KC mice suppresses the UPR. (A) Comparison of the relative protein levels of BiP in normal tissues (Normal, n show as indicated), CCRCC, GBM, HNSCC, LSCC, LUAD, PDAC and UCEC (n show as indicated) in the CPTAC proteomics database. *: *P*<0.05, **: *P*<0.01, ***: *P*<0.001, ns, not significant (unpaired Student’s two-tailed t-test). (B) Comparison of the relative mRNA levels of *HSPA5*, *EIF2AK3*, *ERN1*, *ATF6* and *ATF4* in normal pancreatic tissues (Normal, n show as indicated) and pancreatic tumors (n show as indicated) in the CPTAC mRNA database. *: *P*<0.05; ***: *P*<0.001 (unpaired Student’s two-tailed t-test). (C) Comparison of the relative protein levels of BiP, PERK and IRE1 in normal pancreatic tissues (Normal, n show as indicated) and pancreatic tumors (n show as indicated) in the CPTAC proteomics database. ***: *P*<0.001 (unpaired Student’s two-tailed t-test). (D) IHC staining images of TRIM21 and BiP in pancreatic cancer and normal tissues from Human Protein Atlas database. (E) Comparison of the IOD/area of TRIM21 and BiP in (D) were subjected to densitometric quantitation in normal pancreatic tissues (Normal) and PDAC from Human Protein Atlas database. **: *P*<0.01, ***: *P*<0.001, ****: *P*<0.0001 (unpaired Student’s two-tailed t-test). (F) Interrogation of CPTAC co-expression network demonstrates negative protein correlation (Pearson’s R<-0.5, *P*<0.001) between TRIM21 and BiP in PDAC cohorts. (G) Protein levels of BiP, ATF6, sXBP1, CHOP, ATF4, PERK, IRE1, EIF2A, p-EIF2A and TRIM21 in WT or KC mice in pancreatic tissues. HSP90 was used as a loading control. (H) Quantification of BiP, ATF6, sXBP1, CHOP, ATF4, PERK, IRE1, EIF2A, p-EIF2A and TRIM21/HSP90 ratio in (G). Data were mean ± SD for 3 mice per genotype. *: *P*<0.05; **: *P*<0.01; ***: *P*<0.001 (one-way ANOVA with Dunnett’s multiple comparisons test). (I) PCR-based genotyping verification of KC and *Trim21*^fl/fl^; KC mice.

## KEY RESOURCES TABLE

**Table 1.**
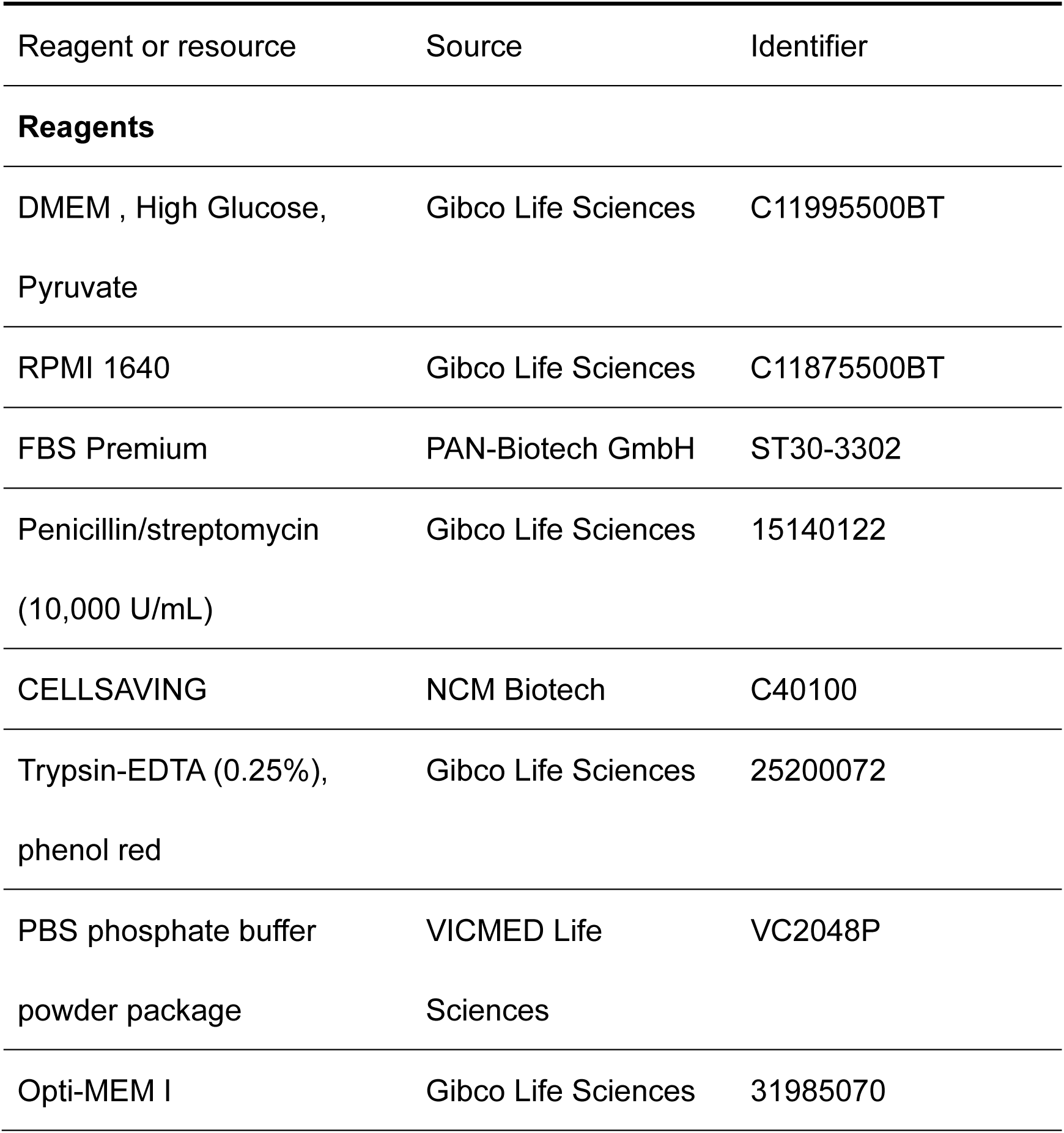
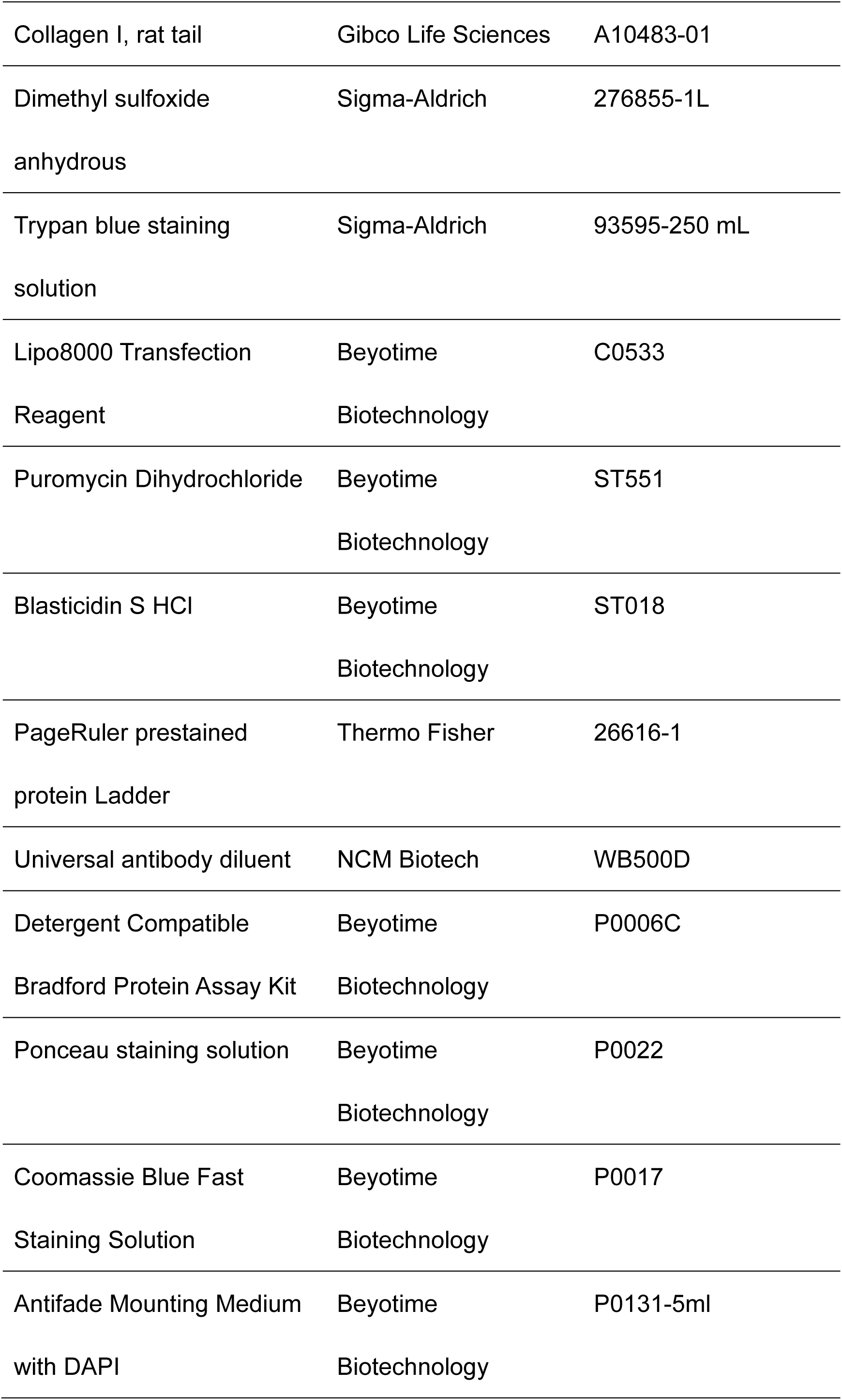
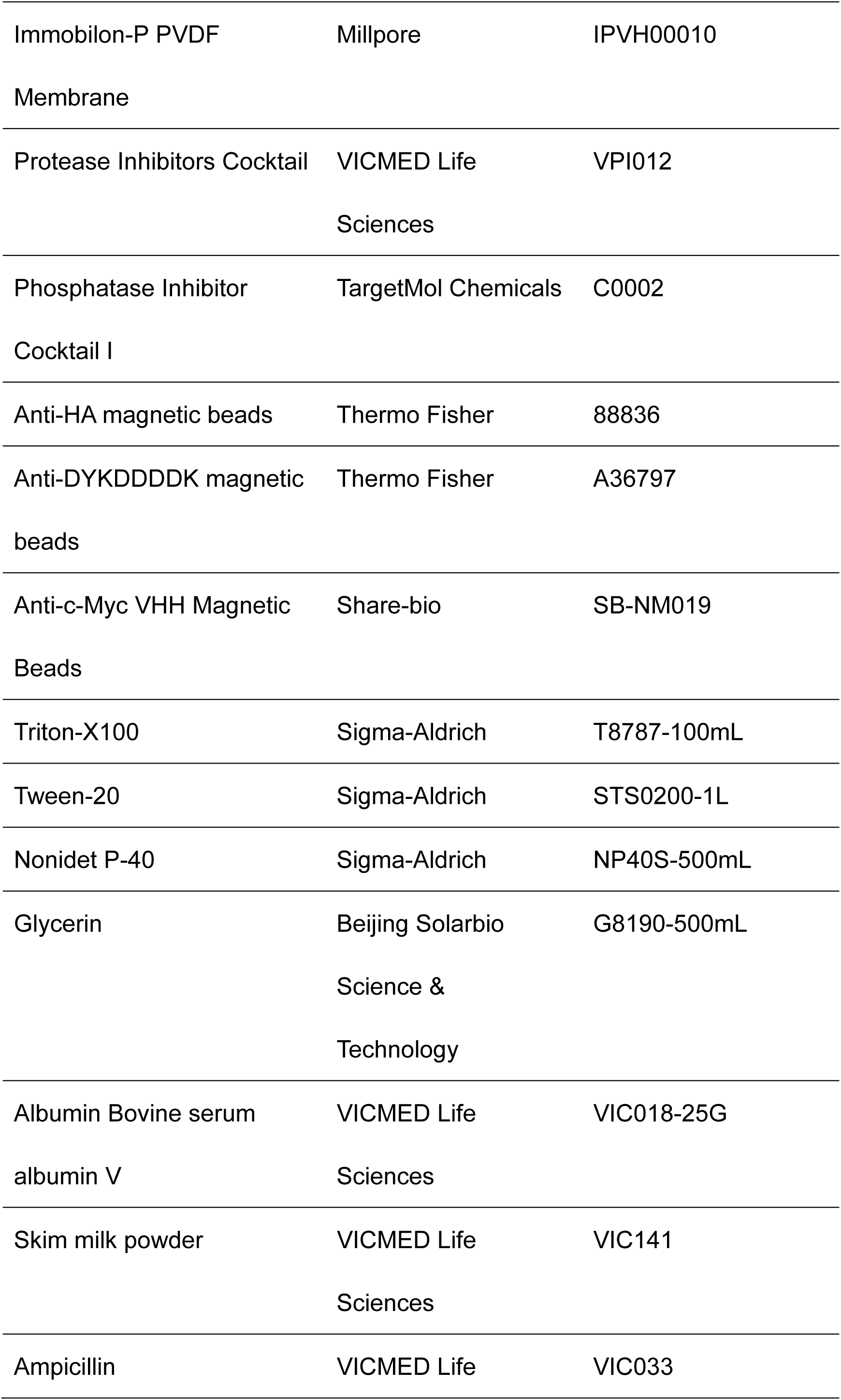
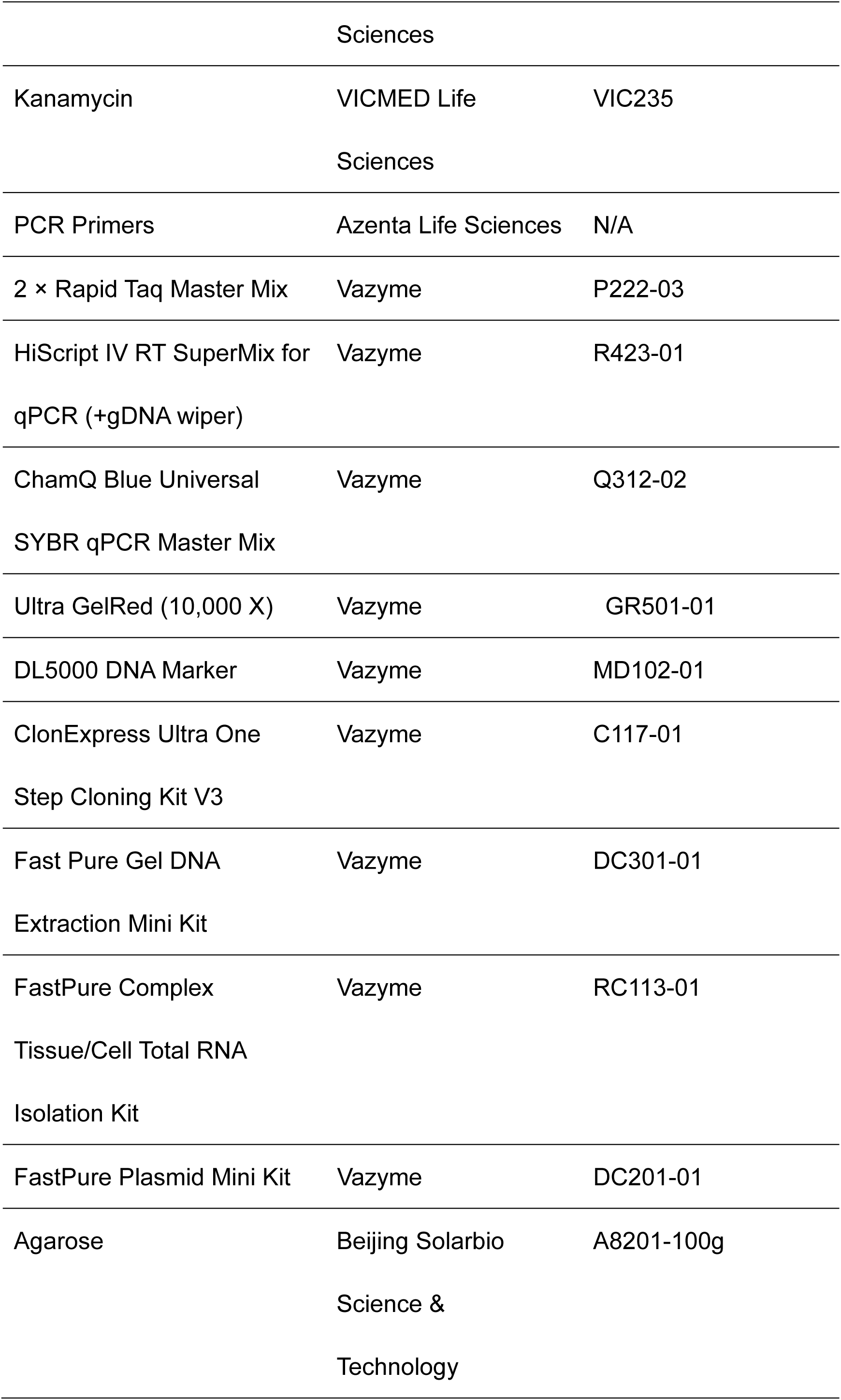
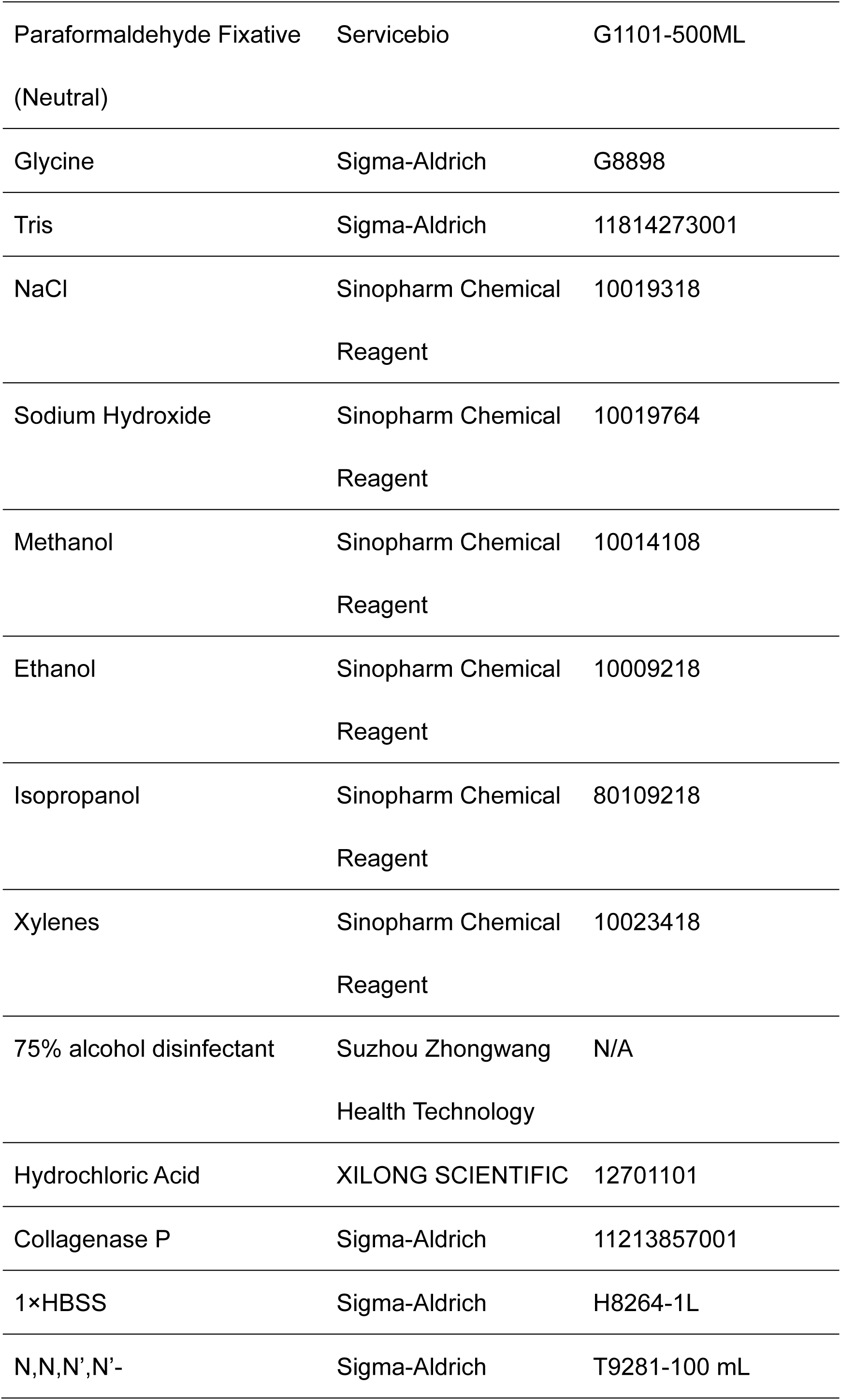
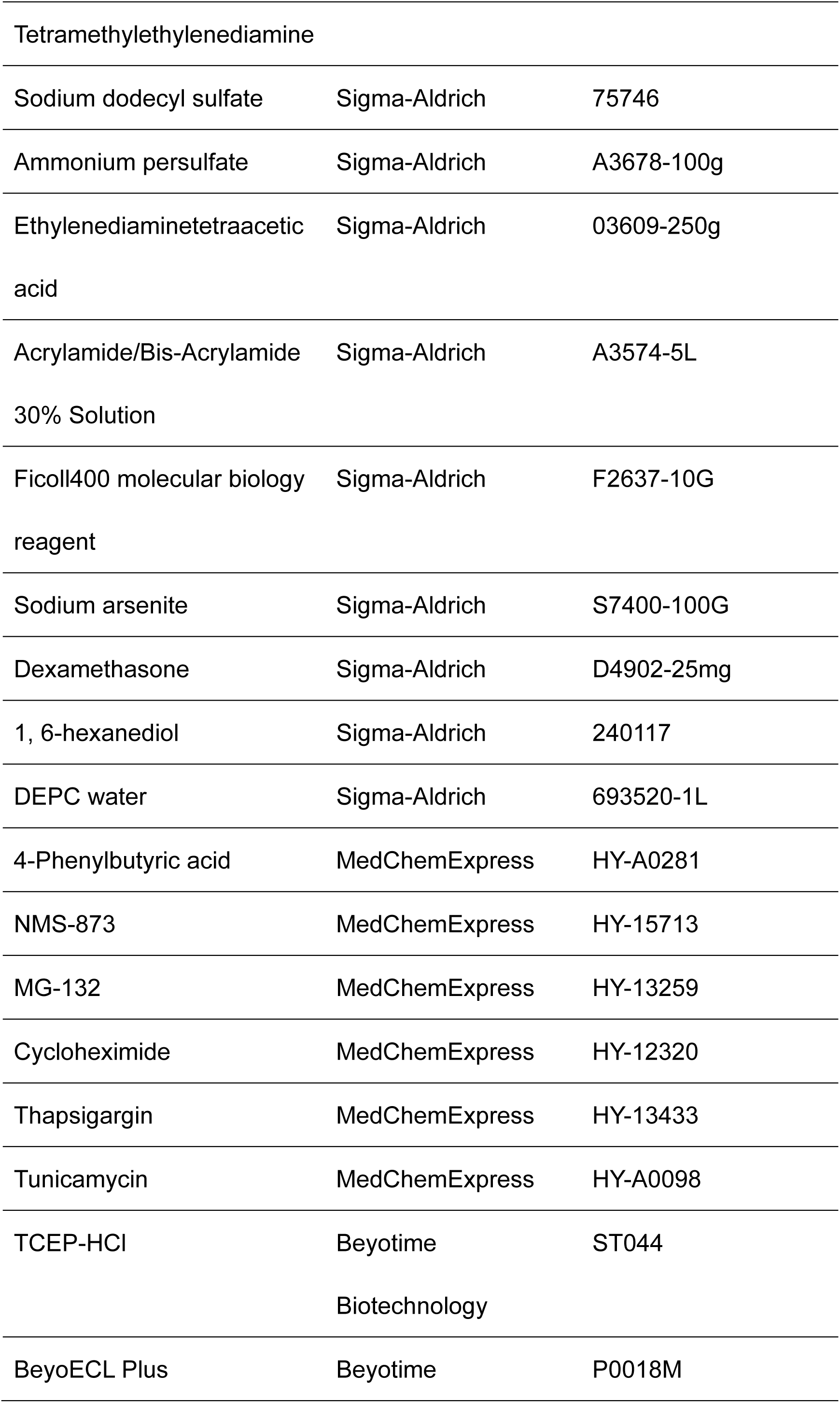
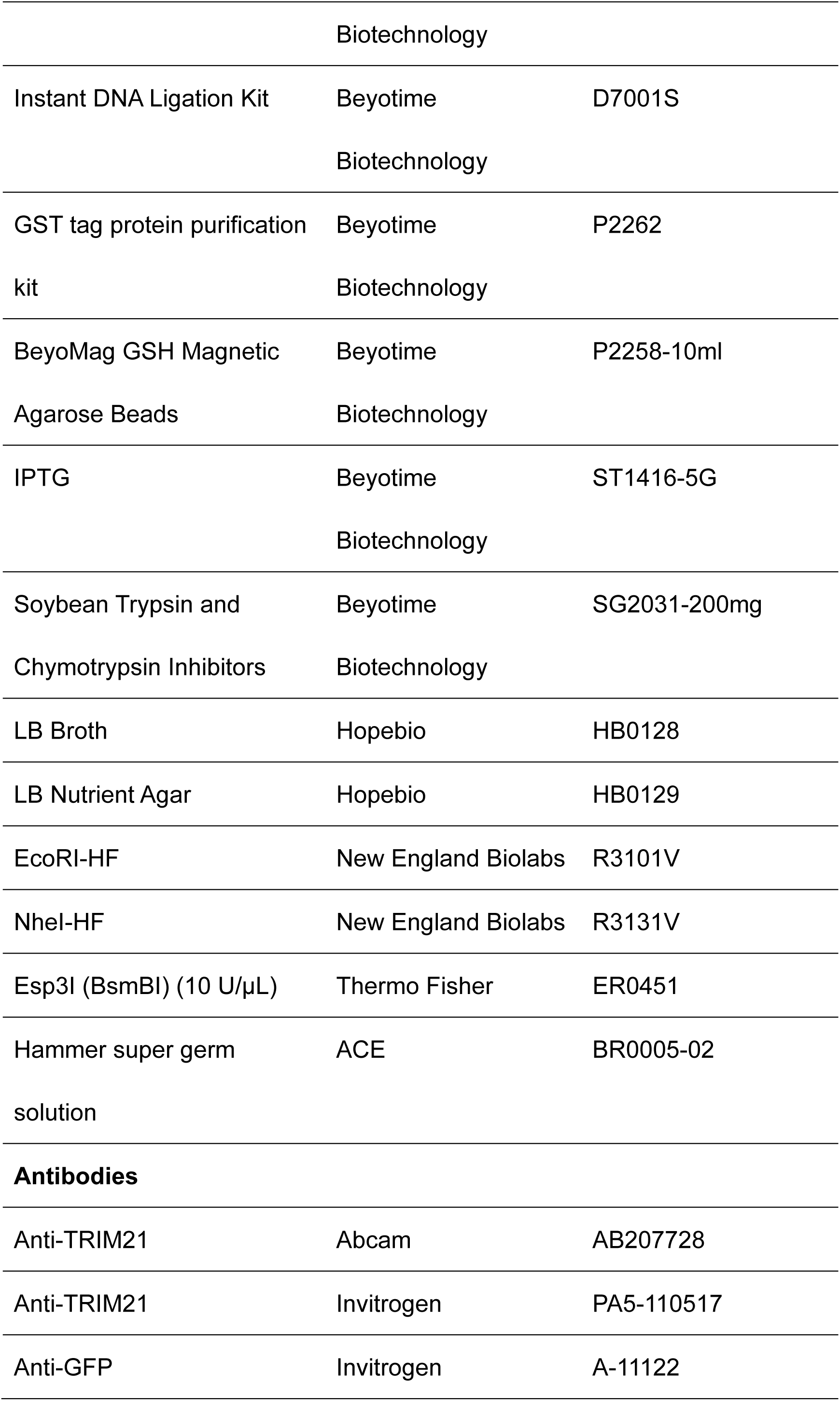
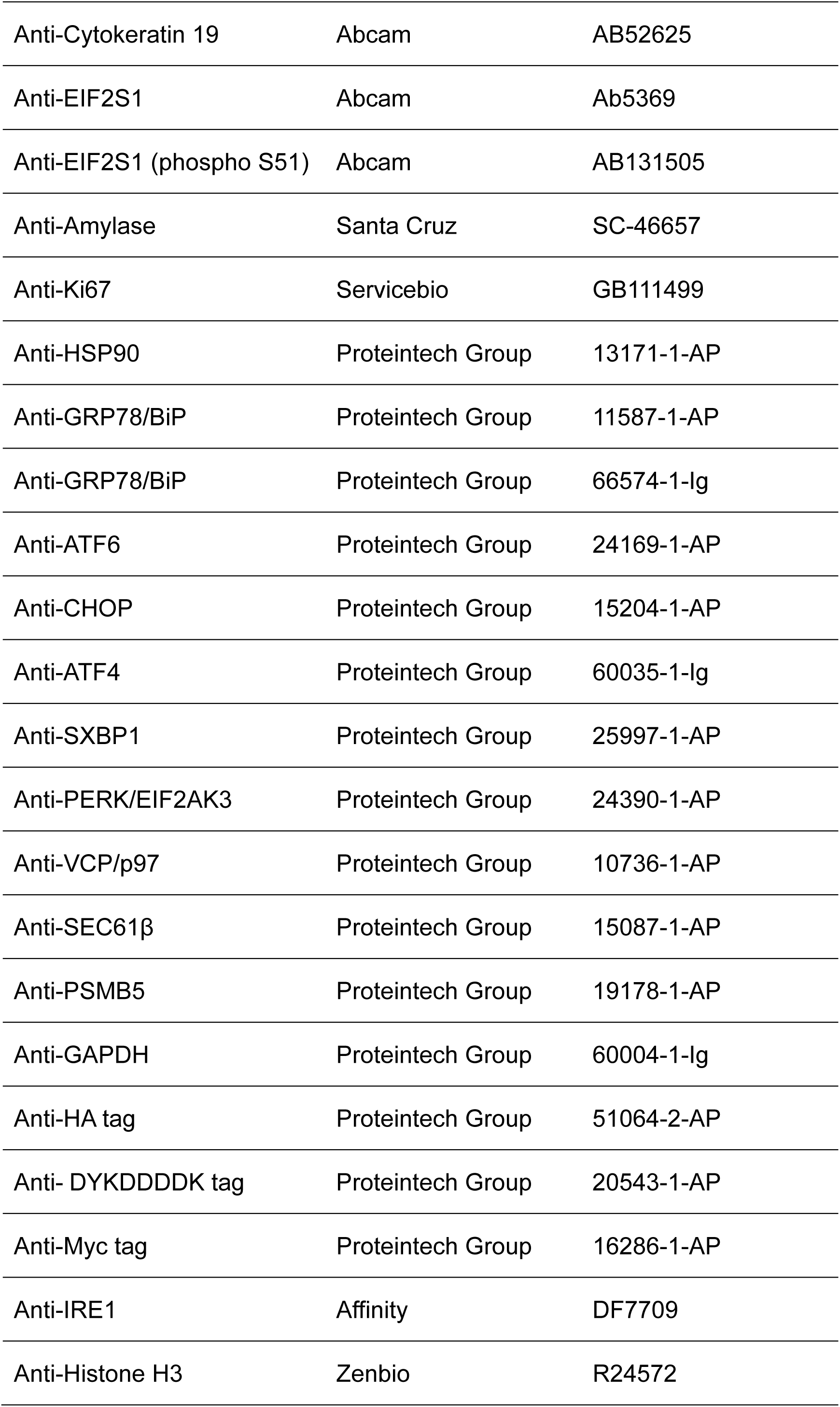
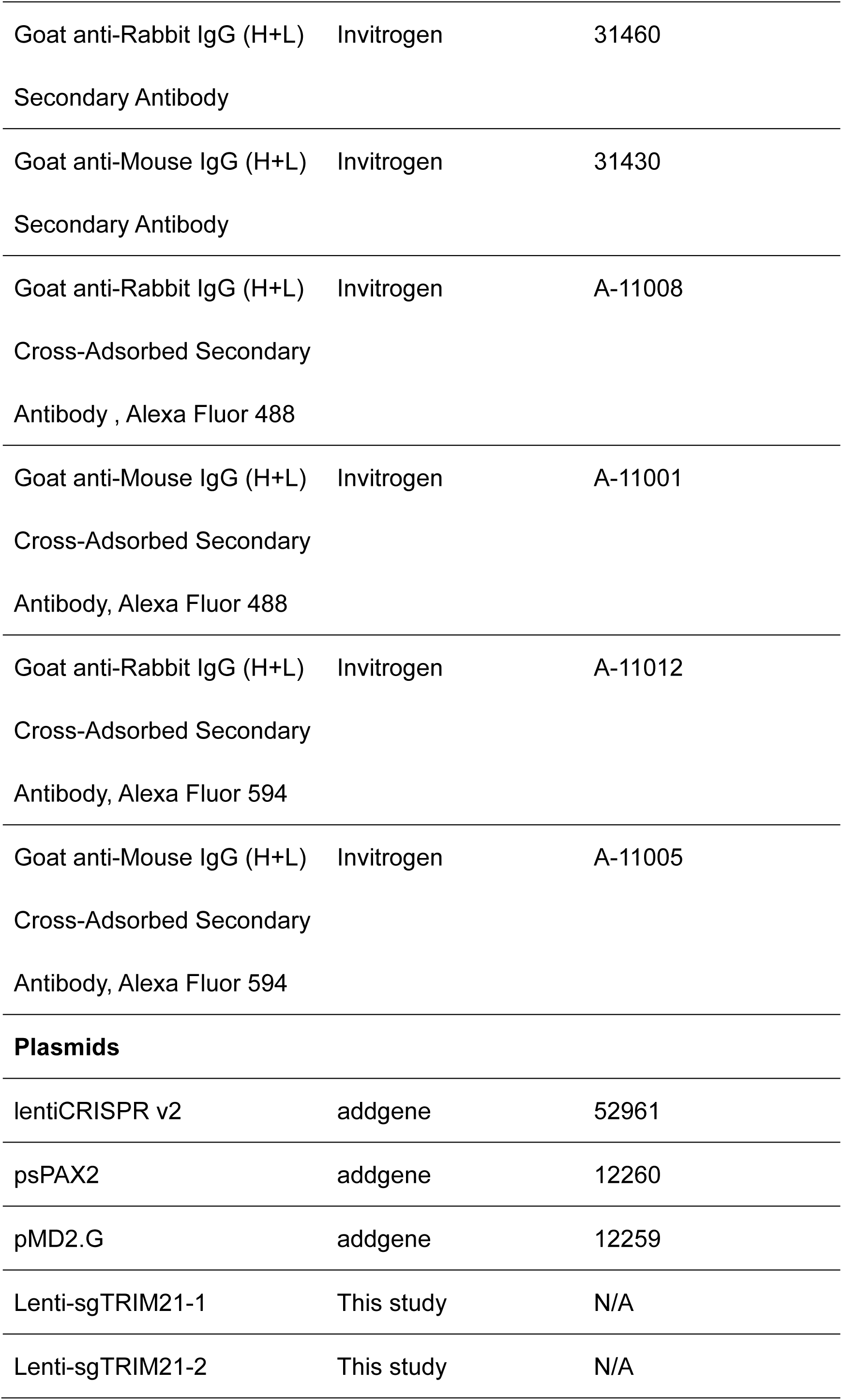
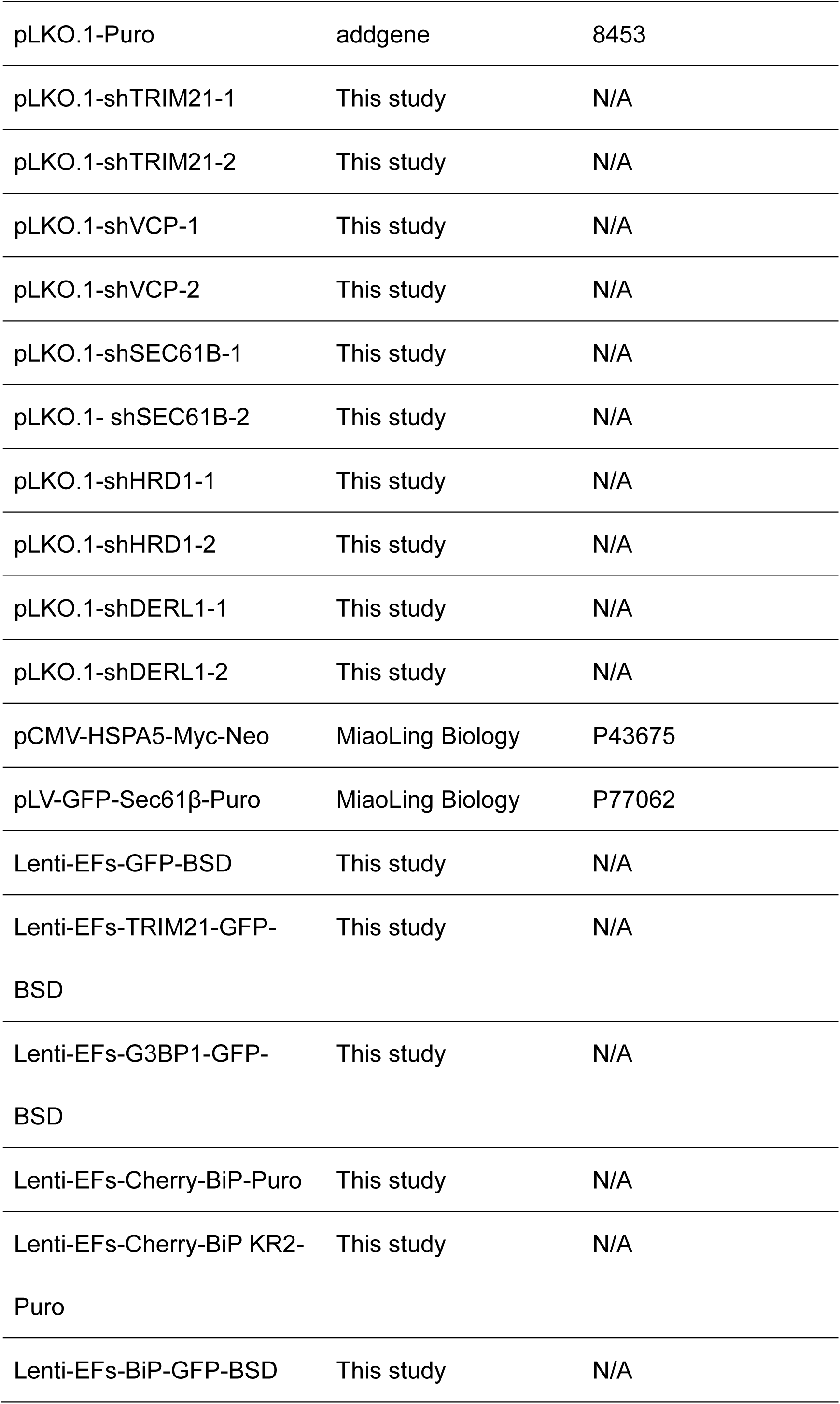
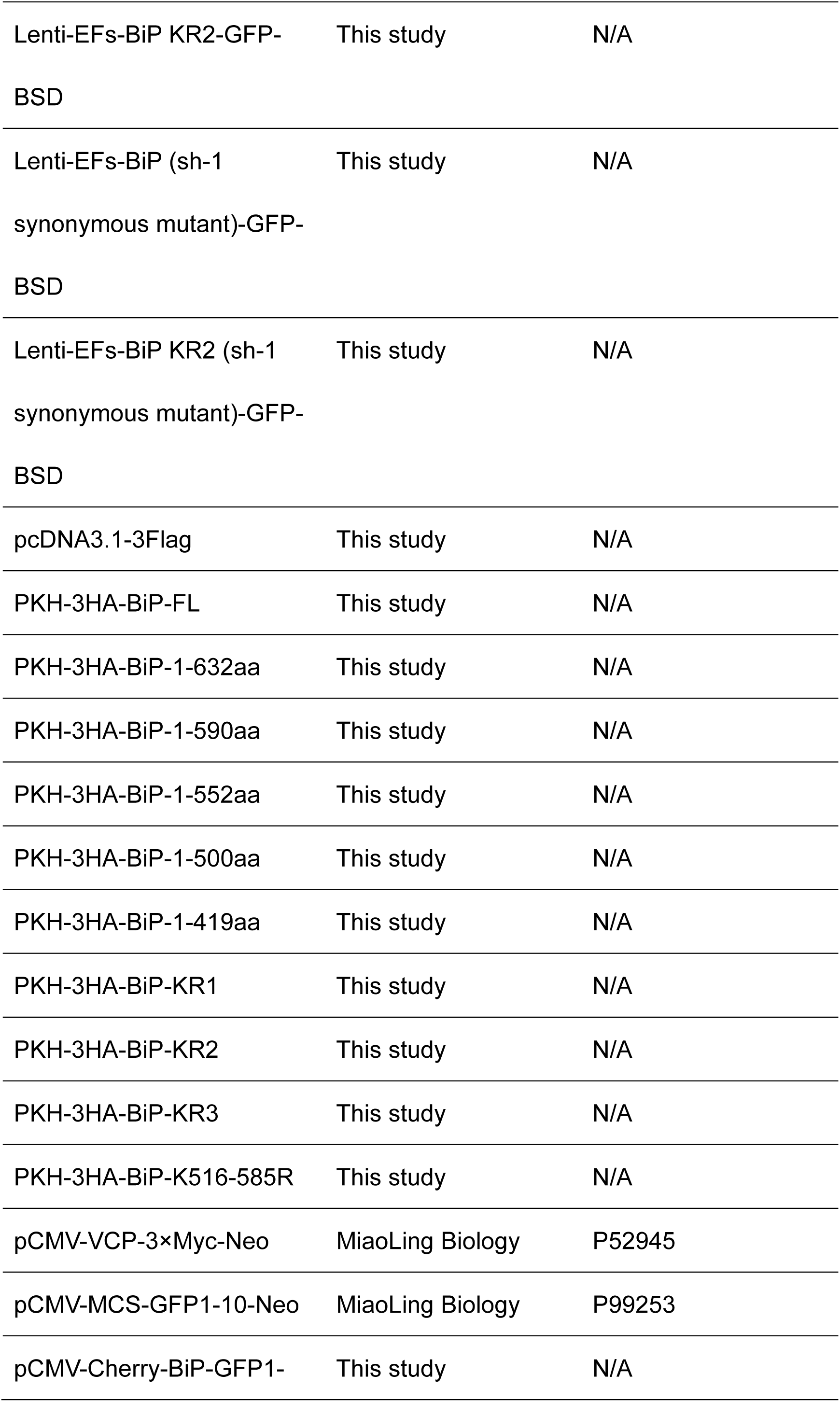
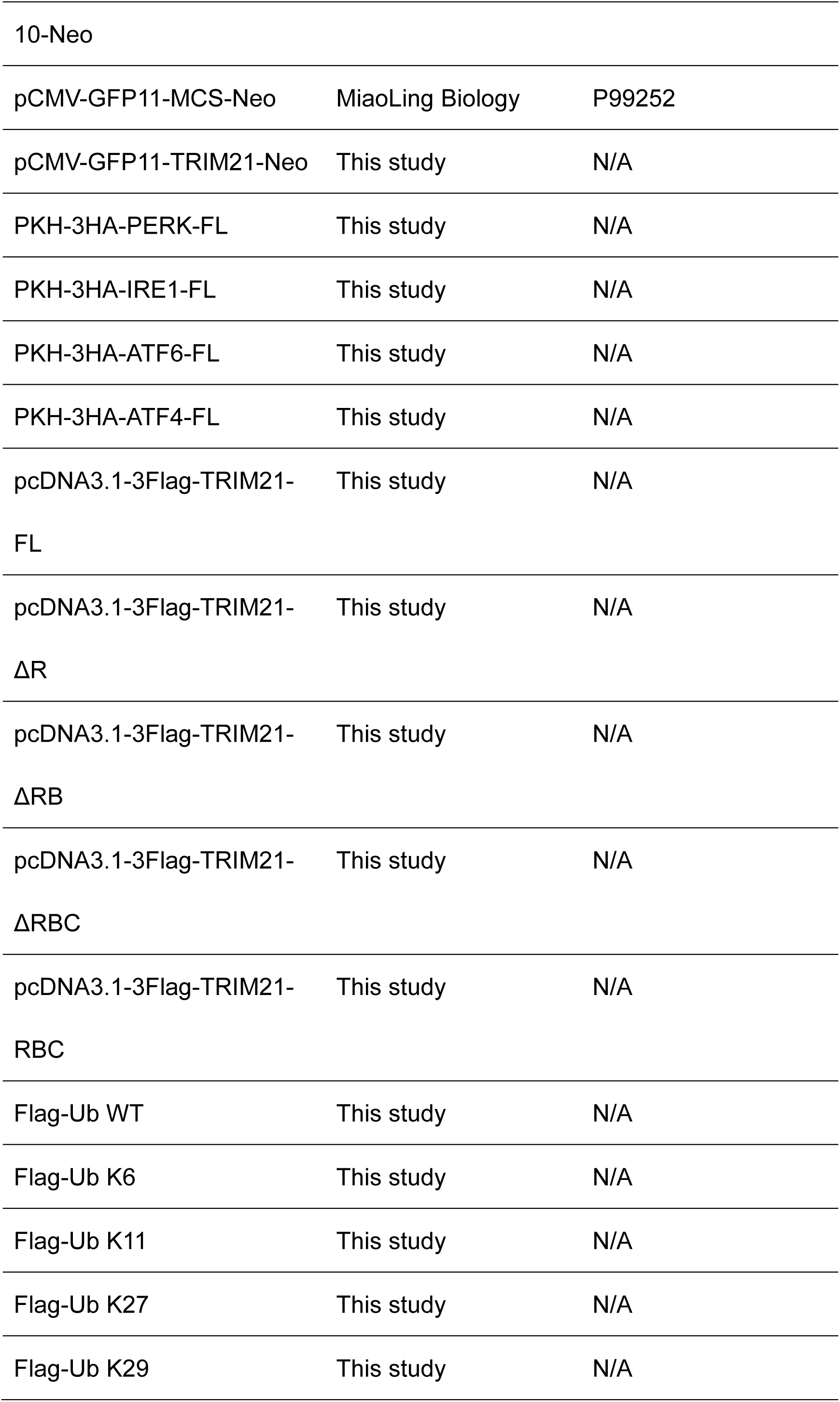
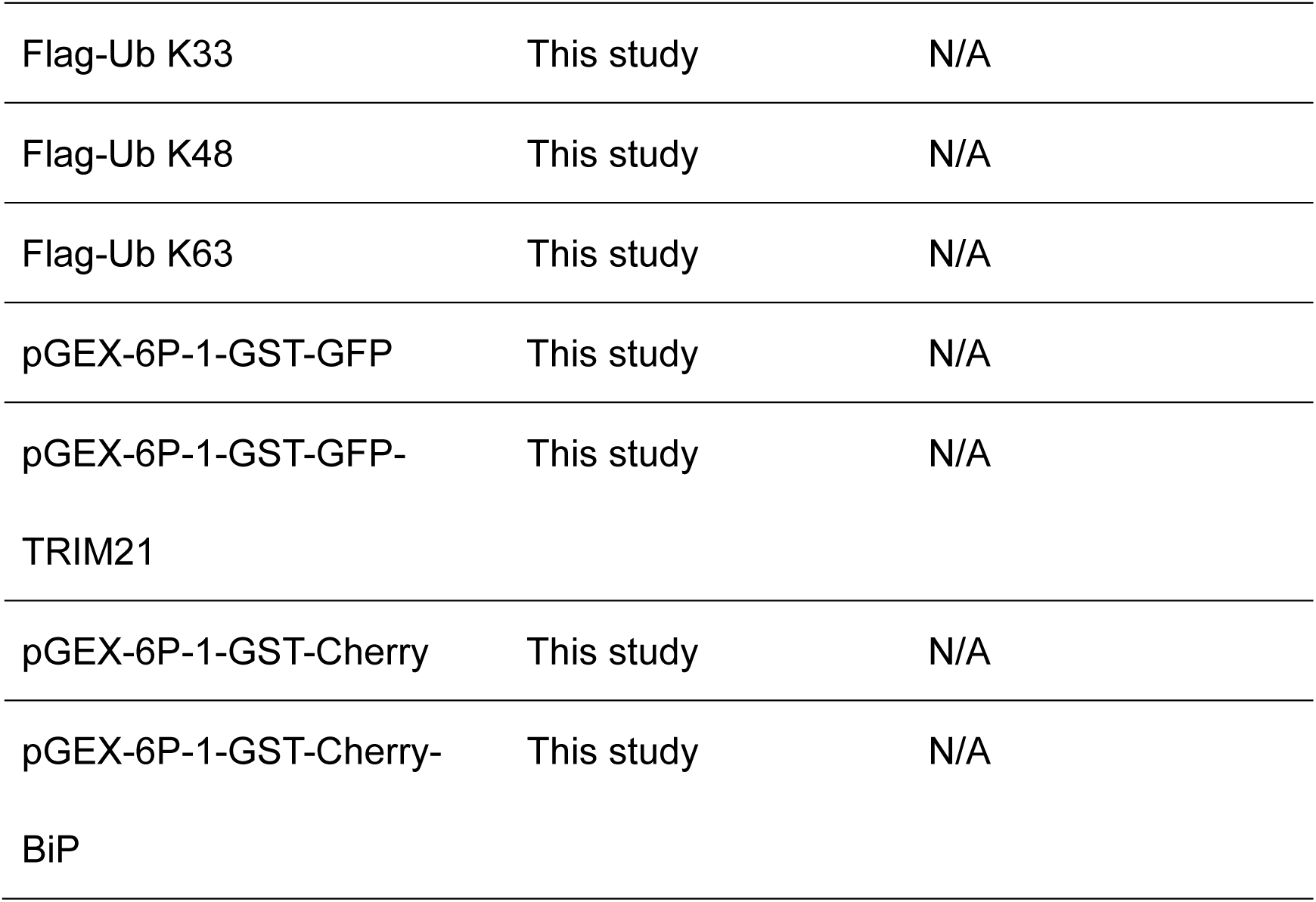
Reagents, antibodies and plasmids.

**Table 2.**
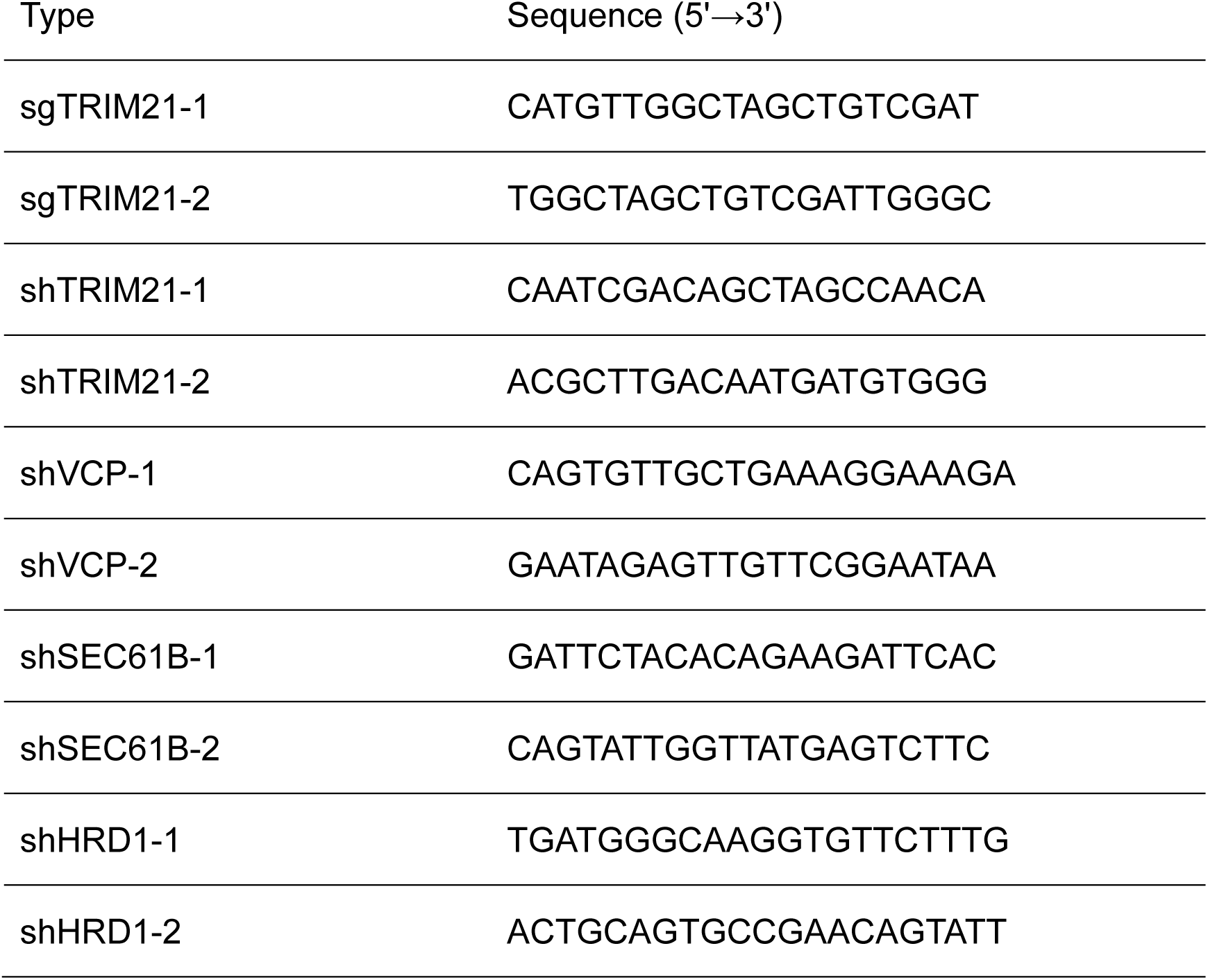
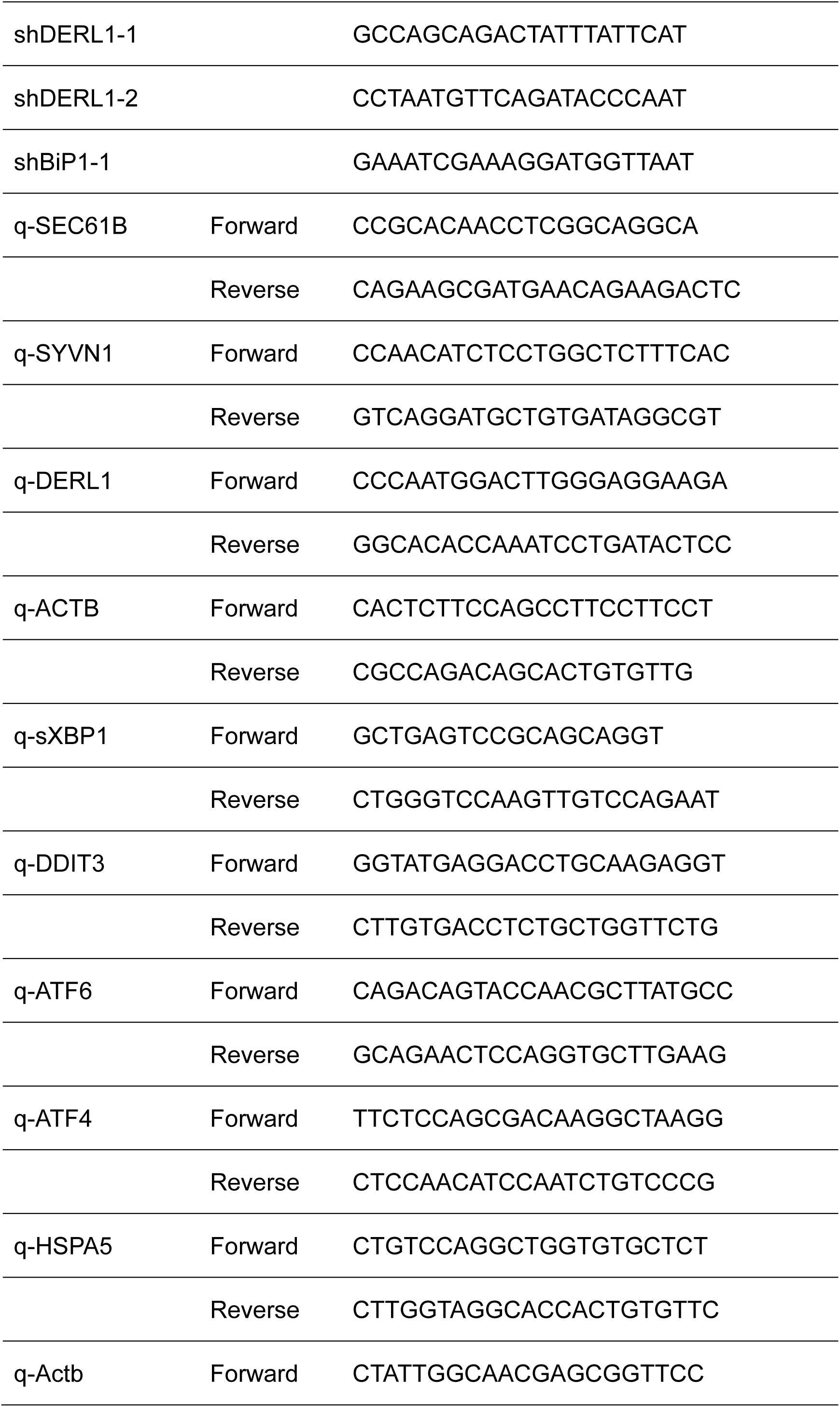
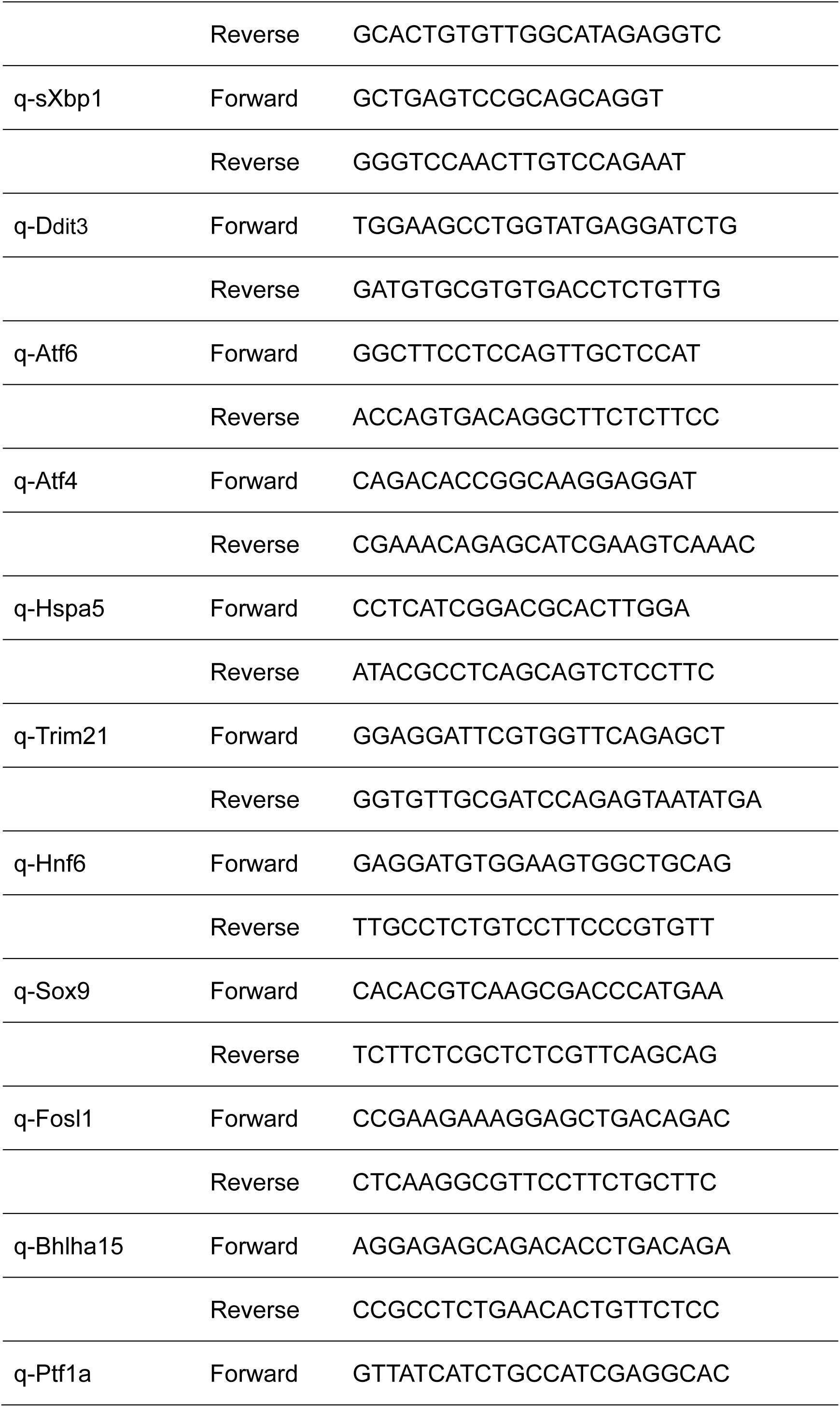
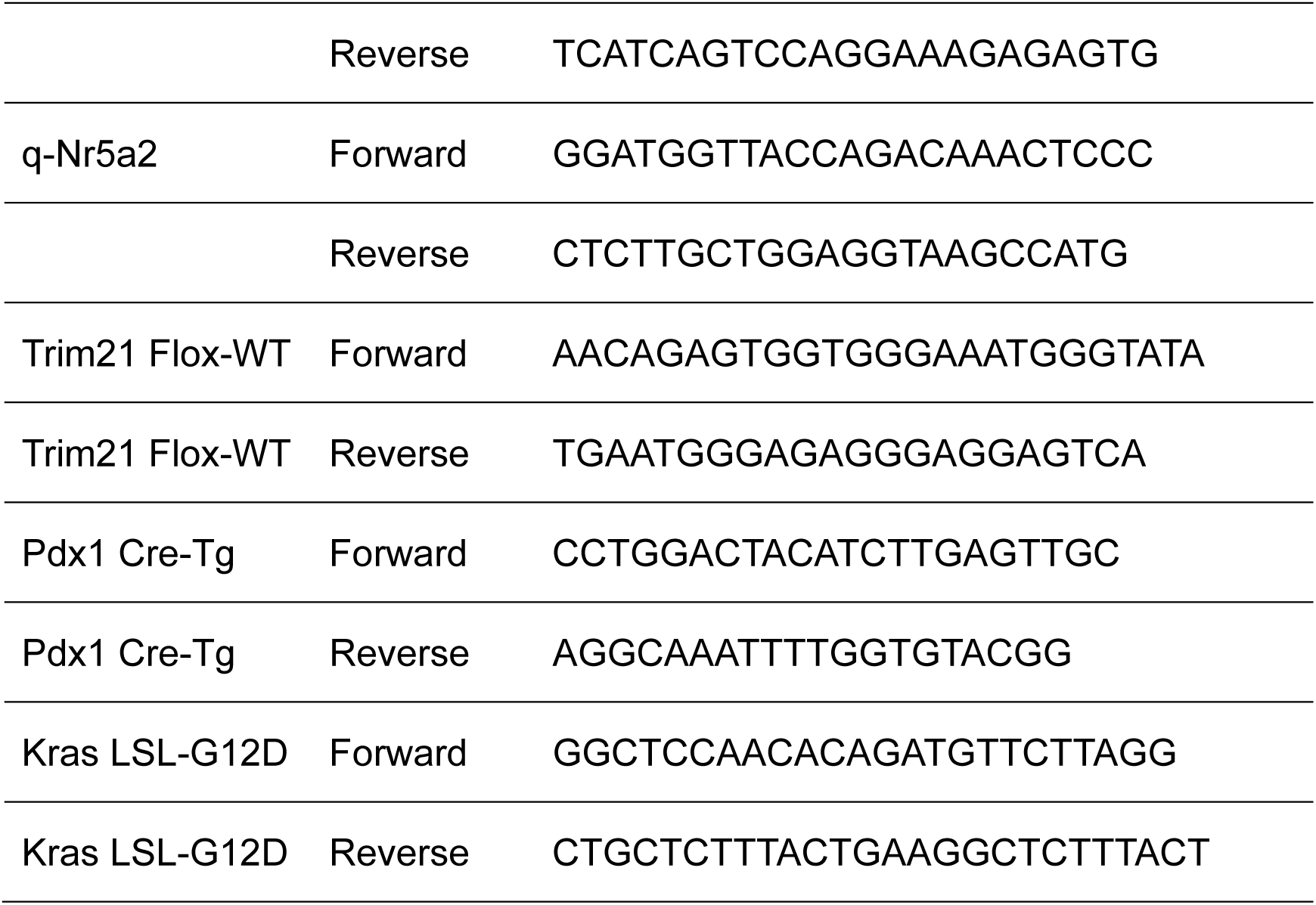
Oligonucleotide sequence.

## EXPERIMENTAL MODEL AND STUDY PARTICIPANT DETAILS

### Cell culture and treatment

Following euthanasia, mice were surface sterilized with 75% ethanol, and pancreatic tissue was dissected and isolated. The tissue was minced and digested with 0.5 mL 2 mg/mL of collagenase P (Sigma) at 37°C with 225 rpm agitation for 25 minutes. The reaction was quenched with ice-cold 1× HBSS containing 5% FBS, followed by three washes with 5 mL of cold 1× HBSS/5% FBS. The cell suspension was filtered through 100 μm strainers (Beyotime), layered over 20 mL of 1× HBSS/30% FBS, and centrifuged at 2000 rpm for 4 mins. Finally, pancreatic acinar cells were cultured in in a type I collagen matrix and maintained in RPMI-1640 medium containing 1% FBS, 1% penicillin-streptomycin, 0.1 mg/mL trypsin inhibitor, and 1 mg/mL dexamethasone. Medium was supplemented on day1 and day3. Tubular ductal structures were counted 2 or 5 days after culturing. HEK-293T (ATCC), Wildtype mouse embryonic fibroblasts (MEF) and PANC-1 (ZQXZBIO) was cultured in DMEM supplemented with 10% FBS, L-Glutamine (2 mmol/L), penicillin (10 Units/mL) and streptomycin (0.1 mg/ml). All cultures were incubated at 37°C in a humidified atmosphere of 5% CO2. Mycoplasma contamination was assessed using the Mycoplasma PCR Detection Kit (Beyotime), and cell line authenticity was confirmed by STR profiling (Wuhan Pricella Biotechnology).

Lentiviral particles were generated in HEK-293T cells by co-transfecting the lentiviral plasmid along with packaging plasmids pMD2.G and pSPAX2 using Lipo8000 Transfection Reagent (Beyotime). 48 hours post-transfection, the culture supernatant containing the lentivirus was harvested, filtered through a 0.45 μm filter, and used to infect target cell lines. Two days after infection, the medium was replaced with puromycin-containing medium for selection. Single-cell clones were isolated by limiting dilution in 96-well plates and expanded for further analysis. Gene overexpression and knockout were validated by Western blot.

### Western blot

Protein was extracted from cellular and murine pancreatic tissue samples using SDS lysis buffer (2% SDS, 62.5 mmol/L Tris pH 6.8, 10% glycerol, 50 mM TCEP). Lysates were boiled for 10 minutes and analyzed by SDS-PAGE on 10% SDS-polyacrylamide gels (Bio-Rad Laboratories) at 130 V for 90 minutes. Subsequently, proteins were transferred to PVDF membranes using a wet transfer system (Bio-Rad Laboratories) at 90 V for 100 minutes.

Membranes were blocked with 5% skim milk/TBST solutions for 1 hour at room temperature to prevent non-specific binding. They were then incubated with primary antibodies, diluted in a universal antibody diluent, either overnight at 4°C or for 4 hours at room temperature. Following three 5-minutes washes with TBST, the membranes were incubated with horseradish peroxidase (HRP)-conjugated secondary antibodies (Invitrogen) for 1 hour at room temperature. After three additional 15 minutes TBST washes, immunocomplexes were detected using BeyoECL Plus (Beyotime) and imaged with a Tanon-5200 Chemiluminescent Imaging System. Protein band quantification was performed using ImageJ software.

### qPCR

Total RNA was extracted using FastPure Complex Tissue/Cell Total RNA Isolation Kit (Vazyme) according to the manufacturer’s instructions before cDNA synthesis using HiScript IV RT SuperMix for qPCR (+gDNA wiper) (Vazyme). The qPCR reactions were performed using the specified primers with ChamQ Blue Universal SYBR qPCR Master Mix (Vazyme). The relative expression was determined using the ΔΔCt method with normalization against β-actin.

### Immunoprecipitation

24 hours after co-transfection with the plasmids of interest, HEK-293T cells were rinsed with PBS and lysed in IP buffer (50 mmol/L Tris-HCl, pH 8.0, 150 mmol/L NaCl, 0.5 mmol/L EDTA, 0.5% NP-40, 10% glycerol) containing protease inhibitors cocktail for 30 minutes at 4°C with gentle agitation. The lysates were centrifuged at 12,000 rpm for 20 minutes at 4°C. The supernatant was incubated for 4 hours at 4°C with Pierce Anti-DYKDDDDK Magnetic Agarose (Invitrogen) or Pierce Anti-HA Magnetic Beads (Invitrogen). The beads were then washed three times with TBST and boiled with SDS-lysis buffer for 10 minutes. Eluted proteins were finally analyzed by immunoblotting.

### Ubiquitination assays

HEK-293T cells were co-transfected for 24 hours with plasmids encoding Flag-Ub, GFP-TRIM21, and HA-BiP. After 6 hours MG132 treatment, cells were then collected with ice-cold PBS at room temperature and pelleted by centrifugation at 1,000 rpm for 5 minutes. After supernatant removal, the cell pellet was lysed in 2% SDS. Lysates were subjected to immunoprecipitation using Pierce Anti-DYKDDDDK Magnetic Agarose (Invitrogen) according to the manufacturer’s instructions. Bound proteins were boiled with SDS-lysis buffer for 10 minutes.

### FRAP assay

PANC-1 cells stably overexpressing BiP-WT-GFP or BiP-KR2-GFP protein through lentiviral transduction were seeded into confocal imaging dishes. FRAP was carried out with a Leica STELLARIS 5 microscope, and the measurements involved five pre-bleaching frames, three flashes of bleaching (100% of laser power at 488 nm), and sixty postbleaching frames (recovery was recorded with an interval of 1.7 s). Data were normalized to the average prebleach fluorescence intensities.

### GST protein expression, purification and in vitro fluorescence-based droplet formation assay

GST-tagged GFP, GFP-TRIM21, Cherry and Cherry-BiP were expressed in T7 Express lysY competent cells by induction with 0.1 mM IPTG for 12 hours at 16°C and affinity purified using glutathione Sepharose beads (Beyotime). After elution using glutathione, the purified GST proteins were concentrated by Amicon Ultra-4 Centrifugal Filter (Millipore). The purified protein was then analyzed using a multichannel live cell imaging system after being combined with Ficoll400 molecular biology reagent (Sigma) (to replicate intracellular crowding conditions).

### GST pull-down assay

The purified GST-GFP or GST-GFP TRIM21 proteins were incubated with HA-BiP lysates at 4°C overnight. Subsequently, GST-GFP/GST-GFP TRIM21 was immobilized onto BeyoMag GSH Magnetic Agarose Beads at 4°C for 2 hours, followed by five additional washing steps. For each experimental sample, 100 μL of elution buffer (10 mM GSH in TBS buffer) was applied, and the eluate was collected by magnetic separation. Elution fractions were analyzed by SDS-PAGE, while the input fractions were analyzed Coomassie brilliant blue staining.

### Histology and immunofluorescence

Following euthanasia via CO_2_ inhalation, mice were dissected for immediate collection of pancreatic tissue. A portion of the tissue was homogenized in lysis buffer for protein extraction, while another subset was preserved in RNAlater solution for subsequent RNA isolation. The remaining tissue was fixed overnight at 4°C in 4% paraformaldehyde. After fixation, tissues were dehydrated through a graded ethanol series (50%, 75%, 85%, 95%, and 100%), with each step lasting 1 hour. Tissues were then cleared in xylene (two changes, 10 minutes each) and infiltrated with paraffin (two changes, 1 hour each). Final impregnation was completed by overnight immersion in paraffin. The following day, tissues were embedded in paraffin blocks, and consecutive 4 μm sections were prepared. These sections were sent to Wuhan Servicebio Technology Co., Ltd. for histopathological analysis, which included hematoxylin and eosin (H&E) staining, Sirius Red staining, Alcian Blue staining and Ki67 immunohistochemistry.

All stained sections were digitized using an Olympus VS120 multispectral high-throughput slide scanner. Quantitative analysis was performed with Image-Pro Plus 6.0 software to measure the following parameters: total pancreatic area, regions of acinar-to-ductal metaplasia (ADM), Alcian Blue-positive low-grade pancreatic intraepithelial neoplasia (PanIN), Sirius Red-positive collagen deposition (fibrosis), amylase-positive intact acini, and the percentage of Ki67-positive cells. Fluorescence images were processed and analyzed using ImageJ software.

Tissue sections were washed in xylene three times (5 mins each) and then rehydrated in graded ethanol solutions (100%, 95%, 85% and 75%, 2 mins each). After heat-activated antigen retrieval in Tris-EDTA pH 9.0 buffer, 6% BSA/TBST solution was used to block the sections at room temperature for 30 minutes. Subsequently, they were incubated with primary antibodies overnight at 4°C in a humidified chamber. Following washes with phosphate-buffered saline (PBS), sections were incubated for 1 hour at room temperature with Alexa Fluor 488- or 594-conjugated secondary antibodies (1:1000 dilution), protected from light. After additional PBS washes, sections were mounted in an anti-fade mounting medium containing DAPI (Beyotime Biotechnology). Images were acquired using an epifluorescence or confocal microscopy (Olympus).

For cell-based immunofluorescence, pancreatic carcinoma cells were plated on glass coverslips in 24-well plates. After fixation in ice-cold methanol for 10 minutes and blocking with 3% BSA in TBST for 30 minutes, cells were incubated with primary and secondary antibodies using the same protocol as described previously.

### CHX chase assay

Cells were transfected with indicated plasmids. After transfection for the indicated times (in Figure legends), cells were treated with 100 mg/ml CHX and were collected at each indicated time point for immunoblot analysis.

### Preparation of the detergent-insoluble fraction

Cells were lysed in soluble lysis buffer (20 mM HEPES pH 7.9, 200 mM KCl, 1mM MgCl_2_, 1% Triton X-100, 1 mM EDTA) (49) containing protease inhibitors cocktail for 10 minutes at 4°C with agitation. Five percent of the resulting lysate was saved as total protein. After centrifugation at 16,000 × g (4 °C for 20 min) the supernatant, or the detergent-soluble fraction, was removed. The pellet, or the detergent-insoluble fraction, was washed with lysis buffer and solubilized with SDS lysis buffer.

### Subcellular fractionation

Cells were lysed in cell fractionation buffer (20 mM HEPES pH 7.9, 10 mM KCl, 1.5 mM MgCl_2_, 2 mM EDTA) (29) containing protease inhibitors cocktail for 10 minutes on ice. Five percent of the resulting lysate was saved as total protein. After adding 0.2% NP-40 for 1 minute on ice and centrifugation at 12,000 × g (4 °C for 10 min) the supernatant, or the cytoplasmic fraction, was removed. The pellet, or the nuclear fraction, was washed with cell fractionation buffer and solubilized with SDS lysis buffer.

### Mass spectrometry

Plasmid-mediated Flag-tagged TRIM21 overexpression in PANC-1 cells was achieved through 48-hour transfection under optimal conditions. Three sequential PBS washes preceded cell disruption in IP buffer containing complete EDTA-free protease inhibitor cocktail (MCE). Flag-tagged protein complexes were immune-precipitated with Anti-FLAG beads (Thermo Fisher) following manufacturer’s protocol. Protein lysates were incubated with affinity matrix for 2 hours at 4°C under constant agitation, with subsequent five-stage washing in high-salt IP buffer. Protein denaturation and release from beads was achieved through 5-minutes incubation at 95°C in reducing SDS-PAGE loading buffer. In-solution trypsinization was performed on purified proteins. LC-MS/MS analysis and raw data processing were conducted by Bioprofile Biotechnology using Q Exactive HF-X system.

### Mice experiments

This study utilized male and female C57BL/6J mice. *Kras*-LSL-G12D (Cat. No. NM-KI-190003) and *Trim21*-Flox (Cat. No. NM-CKO-234039) mice were purchased from The Shanghai Model Organisms Center, Inc. The B6.FVB-Tg(*Pdx1*-cre)6Tuv/J mice (Cat. No. 014647) were purchased from The Jackson Laboratory. Through genetic crossing, we generated the following genotypes: *Kras*^G12D/+^;*Pdx1*^Cre/+^ (KC), *Trim21*^fl/fl^; KC. All mice underwent tail DNA PCR genotyping and were housed in specific pathogen-free (SPF) facilities maintained at 23 ± 1°C with 30-70% relative humidity under a 12-hour light/dark cycle with ad libitum feeding. Animals were monitored weekly, and tissues were collected upon meeting humane endpoints (>10% body weight loss within 7 days, unkempt fur, or hunched posture). Mouse ages are detailed in corresponding figure legends. All procedures were approved by the Institutional Animal Care and Use Committee of Xuzhou Medical University (Approval No. 202406T021) and complied with national laboratory animal welfare standards.

### Database analysis

RNA and protein expression profiles of PDAC and normal tissue and correlation analysis were performed by PCAS (41) (https://jingle.shinyapps.io/PCAS/) with CPTAC proteomics database.

### Statistical analysis

Statistical significance was determined using GraphPad Prism 6, using the unpaired Student’s two-tailed t-test for one-way comparisons, or one-way/two-way ANOVA for multiple comparisons. No statistical methods were used to determine the sample sizes. Unless expressly noted, experiments were not randomized, and the investigators were not blinded to group allocations and the assessment of outcomes. Data were presented as the mean ± SD (error bars) from at least 3 independent experiments. *: *P* < 0.05; **: *P* < 0.01; ***: *P* < 0.001; and ****: *P* < 0.0001.

## Notes

### Competing Interest Statement

The authors have declared no competing interest.

